# Distinct mesenchymal cell states mediate prostate cancer progression

**DOI:** 10.1101/2023.03.29.534769

**Authors:** Hubert Pakula, Mohamed Omar, Ryan Carelli, Filippo Pederzoli, Giuseppe Nicolò Fanelli, Tania Pannellini, Lucie Van Emmenis, Silvia Rodrigues, Caroline Fidalgo-Ribeiro, Pier V. Nuzzo, Nicholas J. Brady, Madhavi Jere, Caitlin Unkenholz, Mohammad K. Alexanderani, Francesca Khani, Francisca Nunes de Almeida, Cory Abate-Shen, Matthew B Greenblatt, David S. Rickman, Christopher E. Barbieri, Brian D. Robinson, Luigi Marchionni, Massimo Loda

## Abstract

Alterations in tumor stroma influence prostate cancer progression and metastatic potential. However, the molecular underpinnings of this stromal-epithelial crosstalk are largely unknown. Here, we compare mesenchymal cells from four genetically engineered mouse models (GEMMs) of prostate cancer representing different stages of the disease to their wild-type (WT) counterparts by single-cell RNA sequencing (scRNA-seq) and, ultimately, to human tumors with comparable genotypes. We identified 8 transcriptionally and functionally distinct stromal populations responsible for common and GEMM-specific transcriptional programs. We show that stromal responses are conserved in mouse models and human prostate cancers with the same genomic alterations. We noted striking similarities between the transcriptional profiles of the stroma of murine models of advanced disease and those of of human prostate cancer bone metastases. These profiles were then used to build a robust gene signature that can predict metastatic progression in prostate cancer patients with localized disease and is also associated with progression-free survival independent of Gleason score. Taken together, this offers new evidence that stromal microenvironment mediates prostate cancer progression, further identifying tissue-based biomarkers and potential therapeutic targets of aggressive and metastatic disease.

## Introduction

Prostate cancer (PCa) ranges from indolent to aggressive, castration-resistant prostate cancer (CRPC), which is associated with a poor prognosis^1, 2^. Genetic alterations in epithelial cancer cells however, do not fully explain the different clinical behavior of this malignancy^3, 4^. Previous studies linked stromal gene expression to prostate carcinogenesis and progression^5, 6, 7^, and our group described that stromal transcriptional programs vary in areas surrounding low vs. high Gleason score. Notably, benign stroma is transcriptionally distinct in tumor-vs. non-tumor-bearing specimens, while benign epithelium does not display significant variability^8^. Furthermore, a stromal gene signature enriched in bone remodeling and immune-related pathways, largely overlapping with one derived from human xenografts that eventually metastasized^9^, predicts metastases^8, 9^. Importantly, prior analyses shows that the stroma is composed of heterogenous and diverse cell populations whose roles in mediating the disease progression have yet to be dissected^10^. In addition, whether the stromal microenvironment differs in the presence of diverse epithelial molecular subtypes of PCa remains to be determined. Therefore, genetically engineered mouse models (GEMMs) driven by different mutations and representing the different stages of prostate carcinogenesis can disentangle the complex stromal remodeling in PCa and reveal stroma-epithelial interacrtions in the tumor microenvironment.

The *Tmprss2-ERG* (*T-ERG*) knock-in murine model^11^ displays a mild epithelial phenotype and serves as a model of PCa initiation. The *Nkx3.1creERT2;Pten^f/f^* (*NP*) mice^12, 13^, and the *Tg(ARR2/Pbsn-MYC)7Key* (*Hi-MYC*) GEMMs^14^ represent prostatic intraepithelial neoplasia (PIN) with subsequent invasion. Advanced, aggressive, invasive adenocarcinoma and neuroendocrine prostate cancer (NEPC) is represented by the *Pb-Cre4 ^+/-^;Pten ^f/f^; Rb1 ^f/f^;LSL-MYCN ^+/+^* (*PRN)* model^15, 16^.

To dissect in detail the tumor microenvironment (TME), and specifically the mesenchymal cells associated with the distinct epithelial lesions present in these GEMMs, we generated a comprehensive single-cell transcriptomic (scRNA-seq) compendium of the mouse PCa mesenchyme, identifying novel stromal cell subtypes characterized by distinct underlying expression programs, driven by regulatory transcription factors determining specific signaling pathways. Distinct mesenchymal cell populations were common across all GEMMs and wild-type (WT) mice, while others showed unique phenotypes aligning with specific PCa-drivers. We further investigated communications within the mesenchymal cells and between stromal components and other supporting or inflitrating cell types. The discovered regulons and interaction networks uncovered novel roles of PCa stroma, influencing disease course via interactions with both tumor and immune cells. Importantly, there is conservation of cluster identity as well as spatial tissue architecture from murine models to prostate cancer in patients. Our results reveal for the first time defined mesenchymal cell populations that might have distinct roles in mediating PCa progression.

## Results

### Distinct stromal populations associated with different stages of prostate cancer

The stroma of PCa GEMMs differed significantly from that of WT counterparts. In particular, stromal remodeling with an increase in extracellular matrix (ECM) rich intraglandular areas begins early in PCa carcinogenesis, with a progressive and significant expansion of the stromal compartment, as measured by image analysis, in models displaying PIN/microinvasion, which reaches the peak in the aggressive neuroendocrine cancer PRN model. (**Figure S1A-B**). This finding highlights active remodeling of the stroma during tumor progression, suggesting that mesenchymal cells may change in function and composition during tumorigenesis.

To gain further insights into the composition and the function of the mesenchymal populations responsible for this stromal reaction, we collected scRNA-seq profiles of 43,582 genes from 101,853 cells in 38 mice using pooled single cell suspensions of all lobes of the mouse prostate without *a priori* marker selection (**Table S1**). We excluded cells of epithelial, lymphoid, endothelial, and neural origin with appropriate markers based on the expression of canonical marker gene sets (**Table S2**). Subsequently, we identified fibroblasts and myofibroblasts (see **Methods**) based on existing validated gene sets (**Table S2**), yielding a dataset of 8,574 mesenchymal cells. The number of cells and transcripts from all models are shown in supplementary information and **Table S1**. After correcting for batch effects and reducing dimension using a conditional variational autoencoder (VAE) (see **Methods**), we determined the different stromal cell types across all mouse models. To this end, we constructed a k-nearest neighbor graph in the VAE latent space using Euclidean metric, and clustered with the Leiden algorithm.

This analysis revealed 12 stromal cell populations. Based on an analysis of cluster-cluster covariance and overlapping marker genes, three of these clusters were merged, while an additional cluster was removed as it had <5% of cells in WT mice. This resulted in a final number of eight distinct clusters (referred to as c0-c7) (**Figure 1A**). The distribution of the 8 mesenchymal clusters among the various GEMMs is shown in **Figure 1B-C**. Some clusters were shared by GEMMs and WTs (c0-c2), while others were strongly enriched in particular mutant models (c3-c7) (**Figure 1C-D**).

**Figure 1.**
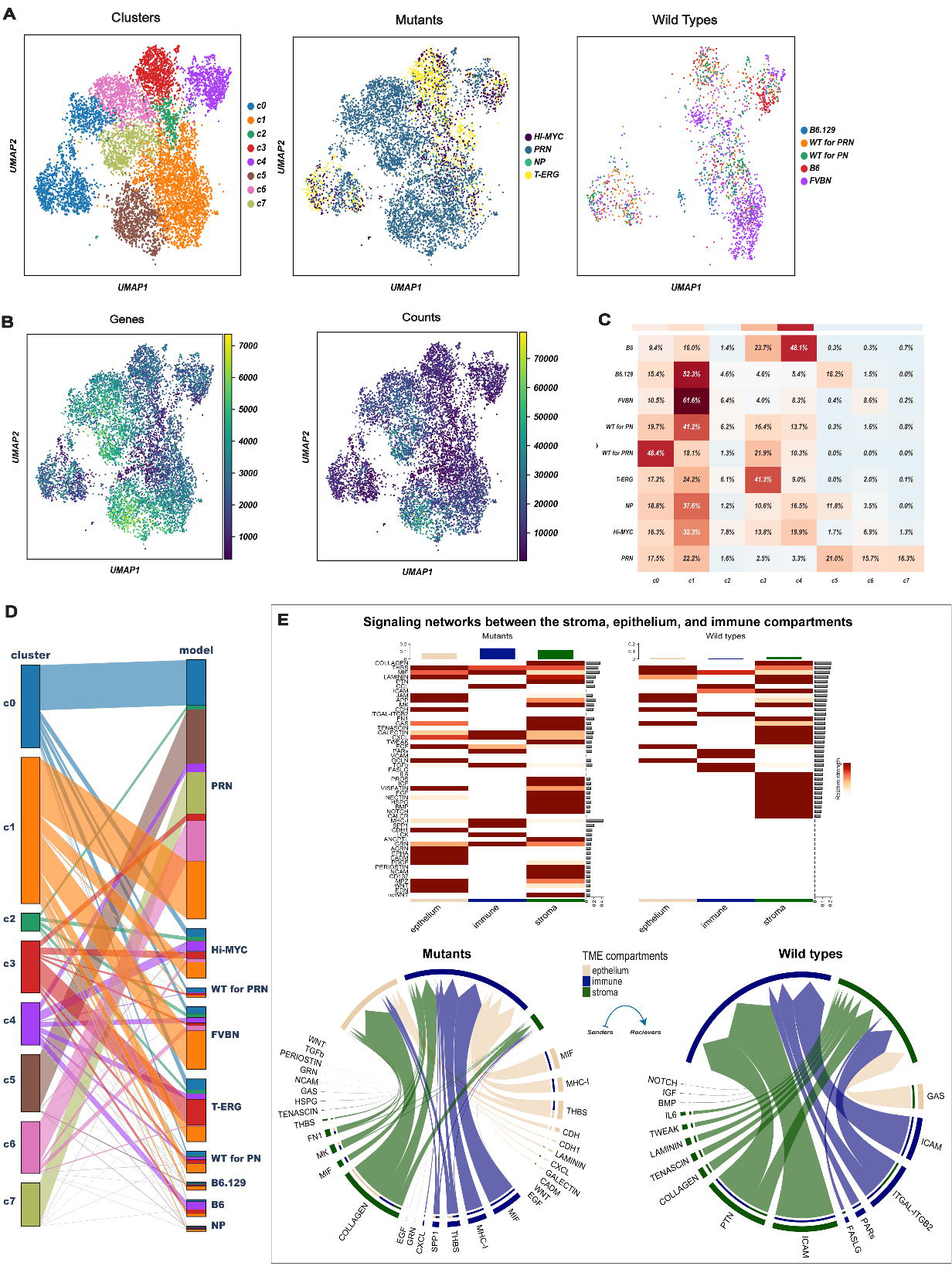
Identification of differentially enriched stromal cell clusters between WT and GEMMs. (A) 8,574 mesenchymal cells visualized by Uniform manifold approximation and projection (UMAP) and colored according to partition assigned by graph-based clustering (left panel) and model of origin (mutant vs. wildtype; middle and right panels). (B) UMAP visualization of the mesenchymal clusters colored by the number of detected genes (left) and Unique Molecular Identifiers (UMIs) (right). (C) Heatmap showing the percentage of the different mesenchymal clusters in each mouse model. Three clusters (c0-c2) represent fibroblast states common to all genotypes, 5 clusters (c3-c7) are specific stromal responses to epithelial mutations. Stroma of two additional wildtype strains (B6 and B6.129) varies in the different backgrounds. (D) Parallel categories plot showing the proportions of mesenchymal clusters (left) across the different mouse models (right). (E) Signaling networks between the stroma, epithelium, and immune compartments. The heatmaps shows the significant outgoing patterns in the mutants (left) and wild types (right). The color bar represents the relative strength of a signaling pathway across cells. The top-colored bar plot shows the total signaling strength of each compartment by summarizing all signaling pathways displayed in the heatmap. The right grey bar plot shows the total signaling strength of a signaling pathway by summarizing all compartments displayed in the heatmap. The chord diagrams display the significant signaling networks between the stroma, epithelium, and immune compartments in mutants (left) and wild types (right). Each sector represents a different compartment, and the size of the inner bars represents the signal strength received by their targets. The heatmap is based on comparing the communication probabilities between mutants and wild types while in the chord diagrams, up- and down-regulated signaling ligand-receptor pairs were identified based on differential gene expression analysis between mutants and wild types. In all cases, we adjusted for the number of cells.

Using ligand-receptor (LR) interaction analysis, we compared both the number and strength of signaling interactions between the stroma, epithelium, and immune compartments in both WT and GEMMs where both were much higher (**Figure 1E**). Specifically, outgoing signaling from the stroma of GEMMs was mediated mainly by the COLLAGEN signaling pathway followed by other pathways that were not active in the WT including WNT, PERIOSTIN, and TGFß (**Figure 1E**). The stromal-epithelium interactions were dominated by THBS, MIF, and WNT signaling pathways, especially in GEMMs (**Figure 1E**).

Since transcription factors can play a role in cell lineage determination, knowledge of driving Gene Regulatory Network (GRN) would improve cluster designations^17, 18^. To this end, we performed cis-regulatory network inference to identify potential regulators (regulons) driving either genotypes or clusters^19^. First, modules of highly correlated genes were identified, then pruned to include only those for which a motif of a shared regulator could explain the correlations. Subsequently, we scored the activity of each regulon in each cell and identified a set of regulons with different activity in the eight mesenchymal clusters **(Figure S2).** We then identified differentially expressed genes (DEGs) in all clusters (**Table S3**) using MAST^20^.

### Common mesenchymal clusters across different mouse models

#### Myofibroblasts and pericytes

Contractile marker genes including *Acta2, Myl9, Myh11* and *Tagln,* and “muscle and smooth muscle cells contraction” by Reactome, were found in the c0 cluster. These cells also highly expressed *Mustn1, Angpt2,* and *Notch3,* suggesting a level of transcriptional complexity greater than previously suggested^21, 22, 23^.

Interestingly, two distinct subpopulations of c0, named subclusters c0.1 and c0.2, were found (**Figure 2A**). Mesenchymal cells from c0.1 expressed myofibroblast marker genes *Rspo3, Nrg2, Hopx,* and *Actg2* (**Figure 2B-C**)^22, 23, 24^ while c0.2 overexpressed pericyte markers (*Rgs5, Mef2c, Vtn, Cygb,* and *Pdgfrb)*. Thus, the c0 cluster is composed of both *bona fide* myofibroblasts and pericytes. Although both sub-clusters were represented in all genotypes, sub-cluster 0.1 (c0.1) was predominantly found in *PRN* and sub-cluster 0.2 (c0.2) was enriched in *NP* (**Figure 2A**). Regulon analysis confirmed the separation of these two sub-clusters (**Figure 2D**).

**Figure 2.**
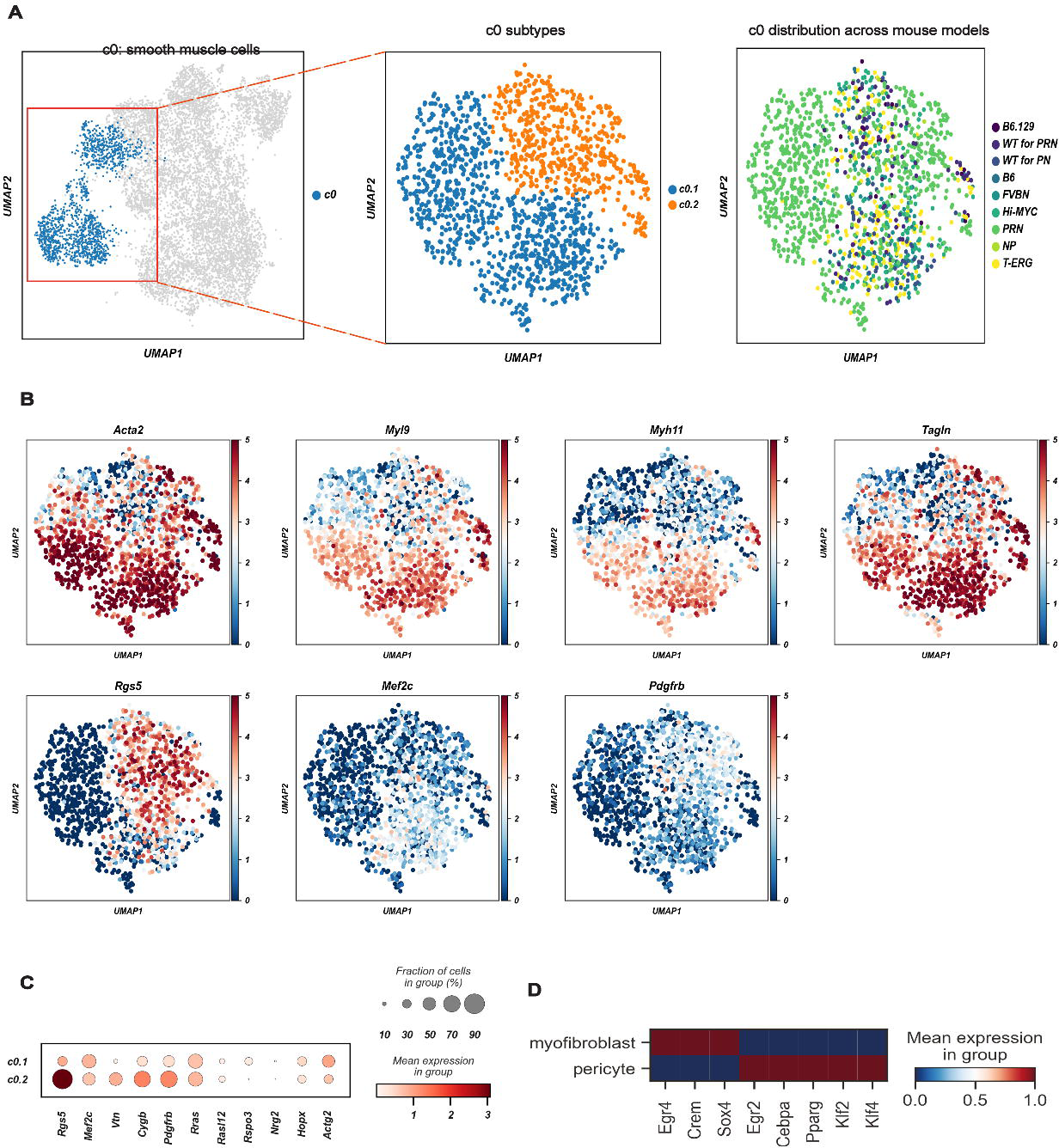
A common cluster of contractile mesenchymal cells encompasses myofibroblasts and pericytes. (A) Canonical myogenic and smooth muscle genes characterize c0 as contractile mesenchymal cells (left panel), but 2 subpopulations (c0.1 and c0.2) may be further subclassified (middle panel). Relative contribution of the different GEMMs and WTs to c0 is shown in the right panel. (B) UMAP projection of c0 cells showing the expression of different myogenic and smooth muscle genes. *Acta2, Myl9, Myh11* and *Tangl* mark myofibroblasts and pericytes, while *Rgs5*, *Mef2c* and *Pdgfrb* distinguish pericytes (c0.2). Color scale is proportional to the expression levels. (C) Dot plot of the expression of genes distinguishing myofibroblasts (c0.1) and pericytes (c0.2). (D) The mean expression of regulons distinguishing myofibroblasts (c0.1) from pericytes (c0.2).

#### Mesenchymal cells with spatially restricted innate immune response genes are conserved across genotypes

The common cluster c1 was characterized by the expression of *Sfrp1* and *Gpx3* (**Figure 3A**) and by major complement system components such as *C3, C7,* and *Cfh*. Validation of the identified c1 markers (Gpx3 and C3) by multiplex immunohistochemistry (mIHC) revealed their substantial enrichment in the stroma surrounding PIN and invasive tumor (**Figure 3B-C**). In addition, c1 showed a unique set of genes overexpressed along with components of immunoregulatory and inflammatory processes (*Ccl11, Cd55, Ptx3,* and *Thbd)* as well as members of the interferon-inducible p200 family of genes (*Ifi204*, *Ifi205,* and *Ifi207)* (**Figure 3A**). Ligand-receptor interaction analysis showed several communication networks from both c0 and c1 to the epithelium mediated mainly by COLLAGEN, LAMININ, and FN1 signaling pathways together with TENASCIN pathway (c1) (**Figure 3D**). Similarly, signaling from c0 and c1 to immune cells in the TME was mediated mainly by the COLLAGEN, THBS, MHC-1, and CXCL pathways together with CCL and COMPLEMENT (c1) pathways (**Figure 3E**). Finally, several regulons, such as *Cebpα* and *Gabpb1,* were identified as well as those governing the inflammatory signaling systems such as *Nfkb1,* along with downstream genes involved in immune activation, which show putative binding sites for these TFs (**Figures 3A** and **S2**).

**Figure 3.**
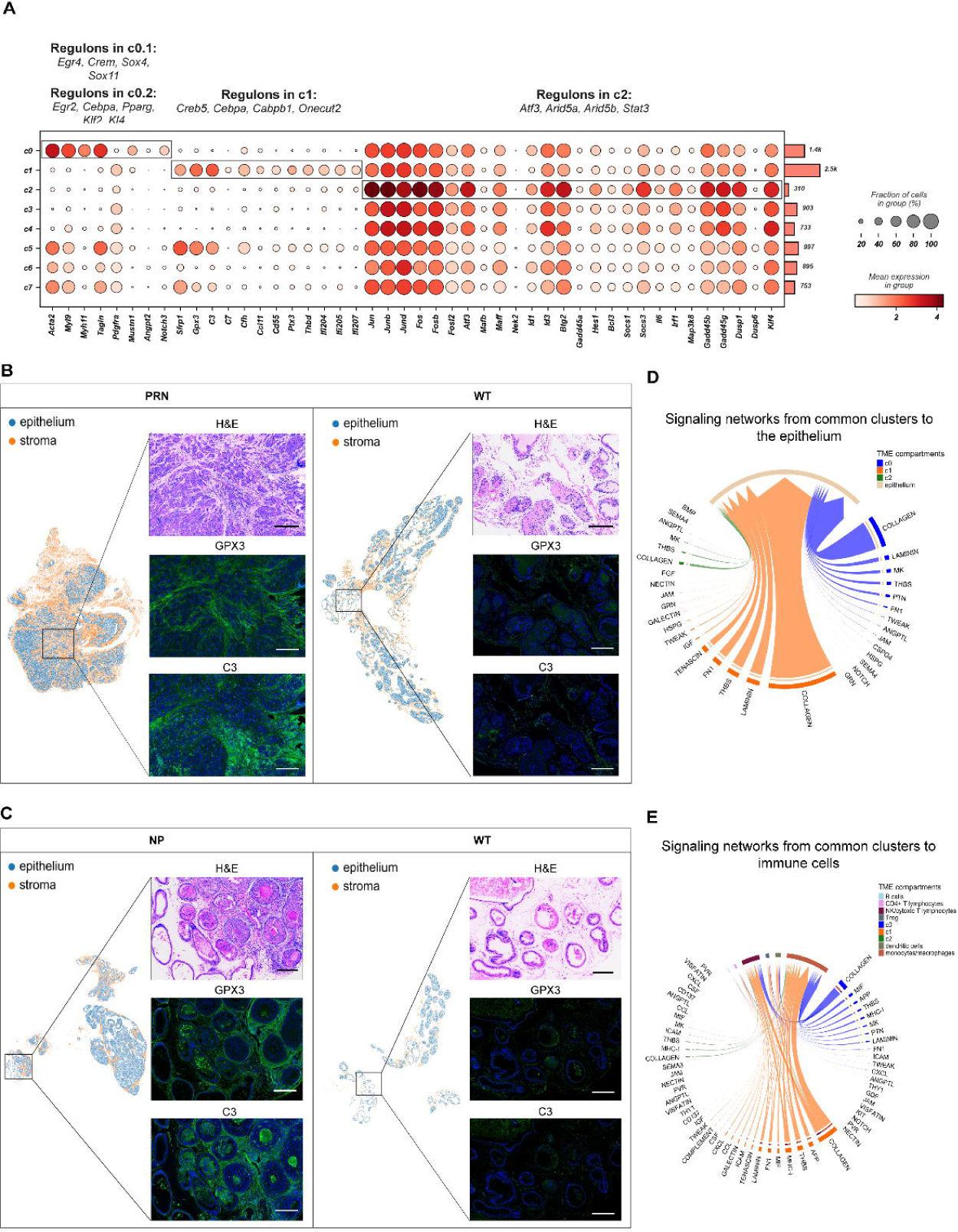
A functional atlas of the mouse prostate cancer mesenchyme. (A) Dot plot showing the mean expression of marker genes for common clusters c0-c2. Boxes indicate the clusters marked by each marker gene set. The total number of cells in each cluster is indicated by the bar plot on the right. Significantly enriched regulons identified by gene regulatory networks are denoted on top of each boxed cluster. (B-C) Representative images of C3 and GPX3 overexpression in tumor desmoplastic stroma in *NP* and *PRN* models (left panels) and matching WTs (right panels). Magnification for all images 200x. Scalebar: 300µm. (D-E) Chord diagrams of significant signaling pathways from the common clusters c0-c2 to the epithelium (D) and immune cells (E). Each sector represents a different cell population, and the size of the inner bars represents the signal strength received by their targets. Communication probsbilities were calculated after adjusting for the number of cells in each cluster.

Mesenchymal cells from c2 were found in all genotypes (**Figure 1C-D**). Components of the c-Jun N-terminal kinase (JNK) pathway were prominently expressed in this population. This was supported by high levels of *Ap-1* components including *Jun*, *JunB, JunD, Fos, FosD, FosB,* and *Fosl2,* activating factors (*Atf3)* (**Figure 3A**). These were concomitant to increased expression of negative regulators of Erk1/2 such as *Dusp1, Dusp6,* and *Klf4* (**Figure 3A)**. GRN analysis revealed candidate TFs regulating MAPK superfamily such as *Atf3, Arid5a* and *Stat3* **(Figures 3A and S2).** Mesenchymal cells in c2 also expressed both negative regulators of the Stat pathway and Stat-induced Stat inhibitors (SSI) (**Figure 3A)**. Interestingly, the expression of SSI family members went along with strong expression of *Il6, Irf1*, which attenuate cytokine signaling. Additionally, similar to c1, c2 interactions with immune cells in the TME were mediated through several signaling pathways including MHC-1, CXCL, and CCL pathways (**Figures 3D-E and S3A-B**).

### GEMM-specific mesenchymal clusters

#### Complex regulation of opposing WNT pathways among stromal clusters

Clusters c3 and c4 were predominantly enriched in *T-ERG, Hi-MYC,* and *NP* models. They expressed core components of the Wnt pathway including genes such as *Sfrp2, Wnt5a, Lgr5, Apc,* enhancers (e.g., *Wnt4* and *Wnt6),* negative regulators of Wnt, (*e.g., Notum* and *Wif1)* as well as TFs such as *Ctnnb1, Lef1,* and *Tcf4* (**Figure 4A**). In-situ validation of c3 and c4 markers by multiplex IHC imaging confirmed the expression of SFRP2 and LGR5 in *T-ERG* mesenchyme compared to the WT stroma. Interestingly, pronounced expression of these two Wnt-proteins was also observed in *PRN* mesenchyme (**Figure 4B-C**). Similar to the common clusters, signaling from c3 and c4 to the epithelium and immune cells was mostly mediated by COLLAGEN and LAMININ pathways combined with a prominent activity of the PTN, THBS, and MK pathways (**Figure 4E**). THBS was also the predominant pathway mediating signaling from monocytes/macrophages (mainly through *Thbs1*) to c3 and c4 (mainly through *Sdc4* and *Cd47*) (**Figure S3C**). Importantly, the immune tumor microenvironment of the NP model had a more prominent infiltration of monocytes/macrophages compared to the other models (**Figure S3A**).

**Figure 4.**
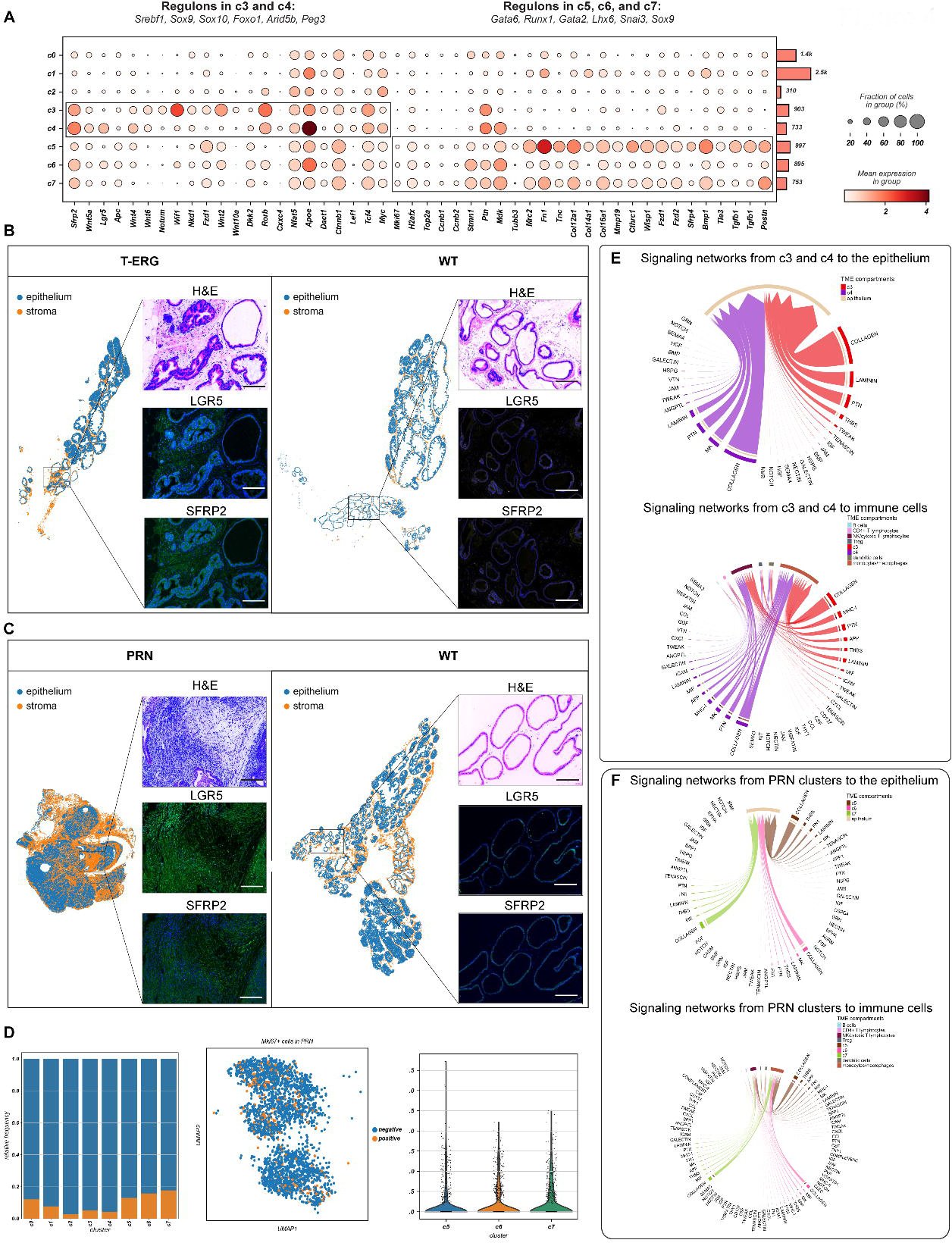
GEMM-specific mesenchymal clusters define complex signaling pathways in the reactive stroma. (A) Dot plot showing the mean expression of marker genes for model-specific clusters c3-c7. Boxes indicate the clusters marked by each marker gene set. The total number of cells in each cluster is indicated by the bar plot on the right. Significantly enriched regulons identified by gene regulatory networks are denoted on top of each boxed cluster. (B-C) Representative images of LGR5 and SFRP2 overexpression in tumor desmoplastic stroma in *T-ERG* and *PRN* models (left panels) and matching wildtypes (right panels). Magnification for all images 200x. Scalebar: 300µm. (D) Bar plots showing the relative frequency of *Mki67*+ mesenchymal cells across all clusters (left), UMAP projections of *Mki67*+ cells in PRN stroma (middle) and violin plots of the expression of *Mki67* in PRN stroma (right). (E) Chord diagrams showing the significant signaling pathways from c3 and c4 to the epithelium and immune cells (upper and lower panels, respectively). (F) Chord diagrams showing the significant signaling pathways from the PRN associated clusters (c5-c7) to the epithelium and immune cells (upper and lower panels, respectively). Communication probsbilities were calculated after adjusting for the number of cells in each cluster.

Several signaling networks between c3, c4 and other stromal cells in the TME especially the PRN clusters (c5-c7) were identified (**Figure S4A**). The WNT and ncWNT signaling pathways in particular were predominantly involved in mediating signaling from c3 and c4 (expressing several WNT ligands like *Wnt5a*, *Wnt2*, and *Wnt4*) to the PRN clusters which expressed several WNT receptors like *Fzd1* and *Fzd2* (**Figure S4B**).

Although both c3 and c4 had similar transcriptional and functional profiles, GRN analysis identified several candidate TFs underlying gene expression differences between the two clusters. For instance, while *Wnt-*stimulatory TFs, including *Sox9* and *Sox10,* were found in c3, *Wnt*-repressive TFs such as *Foxo1* and *Peg3* were enriched in c4 (**Figures 4A and S2**). Overall, these results suggest that the Wnt pathway plays an important yet very complex role in these two clusters.

#### PRN stroma: opposing androgen receptor (Ar) and periostin expression

Cells belonging to clusters c5-c7 were associated with the NEPC mouse models, *PRN*^15, 16^. Generally, cells in these clusters expressed cell cycle and DNA repair-related genes, neuronal markers, a unique repertoire of collagen genes, Tgfβ activation, and again Wnt signaling. Specifically, c5 and 7c expressed high levels of the proliferative markers *Mki67*, γ*H2ax, Top2a*, *Ccnb1*, and *Ccnb2* (**Figure 4A-D**). They also showed high expression of *Cthrc1*, several downstream targets of the Wnt signaling pathway including *Wisp1* and *Ctnnb1*, Wnt receptors such as *Fzd1, Fzd2,* and *Lgr5*, as well as Wnt-secreted decoy receptors *Sfrp4* and *Sfrp2* (**Figure 4A)**. Compared to the other clusters, c5 also expressed the neuronal marker *Tubb3* **(Figure 4A).** The complex stromal response in the PRN mouse model was also highlighted by a unique repertoire of upregulated collagen genes, such as *Col12a1, Col14a1, Col16a1,* and metalloproteinase *Mmp19*, suggesting active remodeling in the tumor microenvironment (**Figure 4A**).

Outgoing signals from the PRN clusters to the epithelium and immune cells showed increased activity of the COLLAGEN, THBS, FN1, MHC-1, CCL, and CXCL pathways (**Figure 4F**). On the other hand, the PRN mesenchyme showed high frequeny and strength of incoming signals from monocytes/macrophages mainly through the THBS and SPP1 pathways and from Tregs through the ITGAL-ITGB2, TGFb, and THY1 pathways (**Figures S3C** and **S4A**). Stromal signaling through the PERIOSTIN pathway in particular was restricted to the PRN mesenchyme with few interactions involving c0 and c1 and no significant interactions involving c3 and c4 (**Figure S4C**).

We again observed activation of adaptive immune responses, such as *C1q* type *a, b* and *c*, in c5-c7 (**Figure S5A**). Finally, stromal NEPC cells highly expressed components of other pathways such as *Nrg1*, *Bmp1*, and *Tgf*β *1, 2, 3* and *Tgf*β-induced *Postn*. Interestingly, high *Postn* expression in c5-c7 was inversely correlated with *Ar* expression, lowest in the PRN model (**Figure 5A**). IF analysis confirmed high POSTN and low AR staining in the stroma (**Figure S6**) especially that adjacent to the invasion front and neuroendocrine foci, while some AR expression was still present around adenocarcinoma foci (**Figure 5B and S6**). In contrast, the highest expression of *Ar* and of its co-regulators *Srebf1*, *Foxo1*, *Arid5b*, *Gata3* and *Creb5* was instead found in c3 and c4, predominantly represented in *T-ERG* and *Hi-MYC* models (**Figure 5A**).

**Figure 5.**
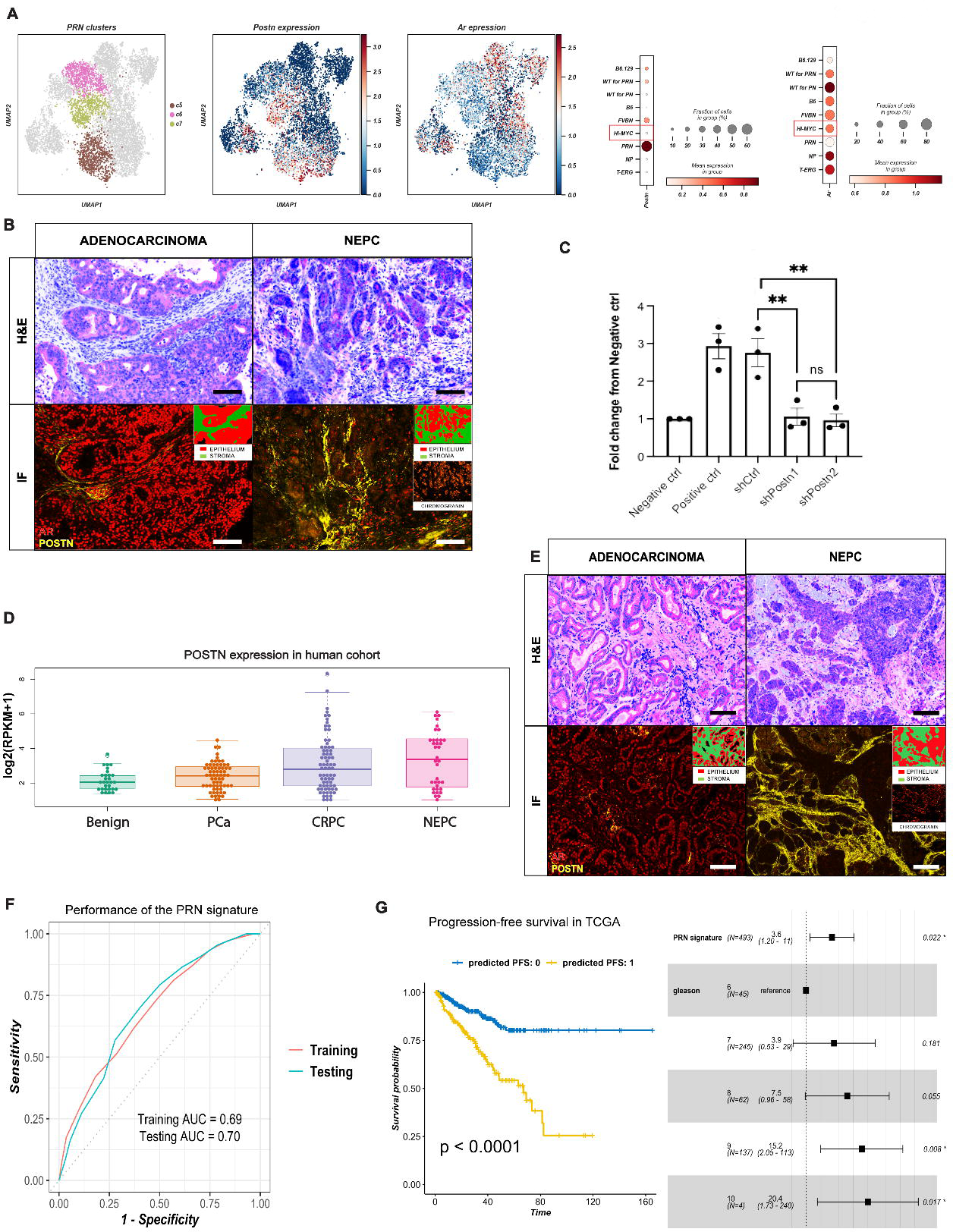
Mesenchymal Periostin overexpression is associated with aggressive, neuroendocrine prostate cancer. (A) UMAP projection of *PRN* clusters c5-c7 (left), *Postn* (middle) *and Ar* (right) expression in prostate mesenchyme; and dot plots showing the mean expression of *Postn* and *Ar* in the different mouse models (right). (B) Multiplexed staining for a panel of proteins including POSTN, AR, and Chromogranin in *PRN* model showing high POSTN and low AR expression in stroma adjacent to neuroendocrine prostate cancer (NEPC) foci (right panel), and weak to moderate AR expression around in the stroma surrounding adenocarcinoma foci (left panel). Magnification for all images 200x. Scalebar: 300µm. (C) Quantification of 22rv1 overexpressing MYCN and with Rb1 knock down migration in Boyden chamber transwell assay. (*** p-value<0.001, ****p-value<0.0001, 1-way ANOVA). (D) Boxplots of *POSTN* expression in WCM clinical cohort. PCa: prostate cancer, CRPC: castration-resistant prostate cancer, NEPC: neuroendocrine prostate cancer, RPKM: reads per kilobase million. (E) Multiplexed staining for a panel of proteins including POSTN (yellow), AR (red), and Chromogranin (orange) in human samples, showing high POSTN and weak to moderate AR expression around the stroma surrounding adenocarcinoma foci (left panel), and high POSTN and low AR expression in stroma adjacent to NEPC foci (right panel) Magnification for all images 150x, scalebar: 300µm. NEPC: neuroendocrine prostate cancer. (F) Receiver Operating Characteristics (ROC) curve showing the performance of the PRN signature at predicting metastasis in the training (n=930) and testing (n=309) data. The signature was trained and tested on bulk expression profiles of primary tumor samples derived from prostate cancer patients. AUC: Area Under the ROC Curve. (G) Progression-free survival (PFS) in the TCGA prostate adenocarcinoma cohort (n=439). On the left, Kaplan-Meier survival plot showing the difference in PFS between patients predicted as ‘0’ and ‘1’ using the PRN signature. The x-axis shows the survival time in months. P: p-value using the logrank test. On the right, forest plot for multivariate Cox proportional hazards model showing the hazard ratio and 95% confidence interval for the PRN signature and Gleason grade. (*p-value <0.05).

In order to assess whether *Postn*-positive stroma facilitates invasion, a characteristic of NEPC, a migration assay was utilized. Knockdown of Periostin in fibroblasts induced an over 2-fold decrease of mobility in 22rv1 cells overexpressing MYCN with additional *Rb1* knockdown to mimick the PRN model (**Figure 5C**). In bulk RNA-seq data from a large cohort of well-characterized benign, locally advanced PCa, CRPC, and NEPC samples (https://shinyproxy.eipm-research.org/app/single-gene-expression), *POSTN* expression was significantly increased in a subset of CRPC and most NEPC patients compared to PCa and benign samples (**Figure 5D**). Finally, mIHC in human cases showed an increased expression of POSTN in the stroma of CRPC and more significantly in NEPC compared to normal prostate stroma (**Figure 5E**). Several regulons driving these clusters involved transcription factors that generally define lineage in mesenchymal stem cells including *Gata6, Runx1*^25, 26, 27^, *Gata2*^28^, *Lhx6,* and *Snai3*^29, 30^ (**Figures 4A** and **S2**).

#### The transcriptional profiles of the PRN-derived clusters are predictive of metastatic progression in prostate cancer

We examined the predictive and prognostic relevance of the PRN-derived clusters (c5-c7) using gene expression profiles of primary tumor samples from a large cohort of PCa patients (n=1239). The expression of the top positive and negative markers of the PRN-derived clusters were used as a biological constraint to train a rank-based classifier of PCa metastasis (see Methods). The resulting PRN gene signature consisted of 13 up-and down-regulated gene pairs from the PRN mesenchyme (**Table S4**). In addition to its interpretable decision rules, this signature had a robust and stable performance in both the training (930 samples) and testing (309 samples) sets with an Area under the Receiver Operating Characteristic Curve (AUC) of 0.69 and 0.70, respectively (**Figure 5F**). Finally, we tested the prognostic value of the signature in the TCGA cohort which included 439 primary tumor samples from PCa patients^31, 32^. In this independent cohort, the PRN signature was significantly associated with progression-free survival (PFS)^31^ using Kaplan-Meier survival analysis (logrank p-value <0.0001), even after adjusting for Gleason grade in a multivariate Cox proportional hazards model (HR=3.6, 95% CI=1.2-11, p-value=0.022) (**Figure 5G**). Overall, these results show that this PRN-derived mesenchymal cell clusters are associated with invasiveness and metastatic progression in PCa patients.

### Human mesenchymal clusters in primary and metastatic tumors

#### Projection of the eight mesenchymal clusters to human scRNA-seq data

Using the mouse scRNA-seq data as reference, we mapped the eight stromal clusters to the human scRNA-seq data*^33^.* These included six ERG-positive (6,990 mesenchymal cells) and three ERG-negative (1,638 mesenchymal cells) patients. c3 was the most predominant cluster in the human stromal data (79% of total mesenchymal cells), (**Figure 6A**) a finding attributed to the selection of ERG-positive cases. Notably, both c0 and c1 had transcriptional profiles similar to their murine counterparts, with c0 characterized by myofibroblast features (*ACTA2, MYL9, MYH11, and TAGLN*), while c1 had a high expression of *SFRP1, GPX3, and C3* (**Figure 6B**). In contrast, the three *PRN*/NEPC-associated clusters (c5-c7) were less abundant in human tumors (13% of total mesenchymal cells) compared to mouse specimens (31% of total mesenchymal cells), a finding explained by the absence of NEPC cases in the human primary tumor cohort. Nonetheless, these *PRN*/NEPC-associated clusters showed a high expression of *C1Qs* (**Figure S5B**), suggesting activation of the adaptive immune response. Overall, the transcriptional similarities of the mesenchymal clusters between both the mouse and human data suggest successful mapping between both datasets despite their biological heterogeneity which is still captured by the difference in cell type frequency.

**Figure 6.**
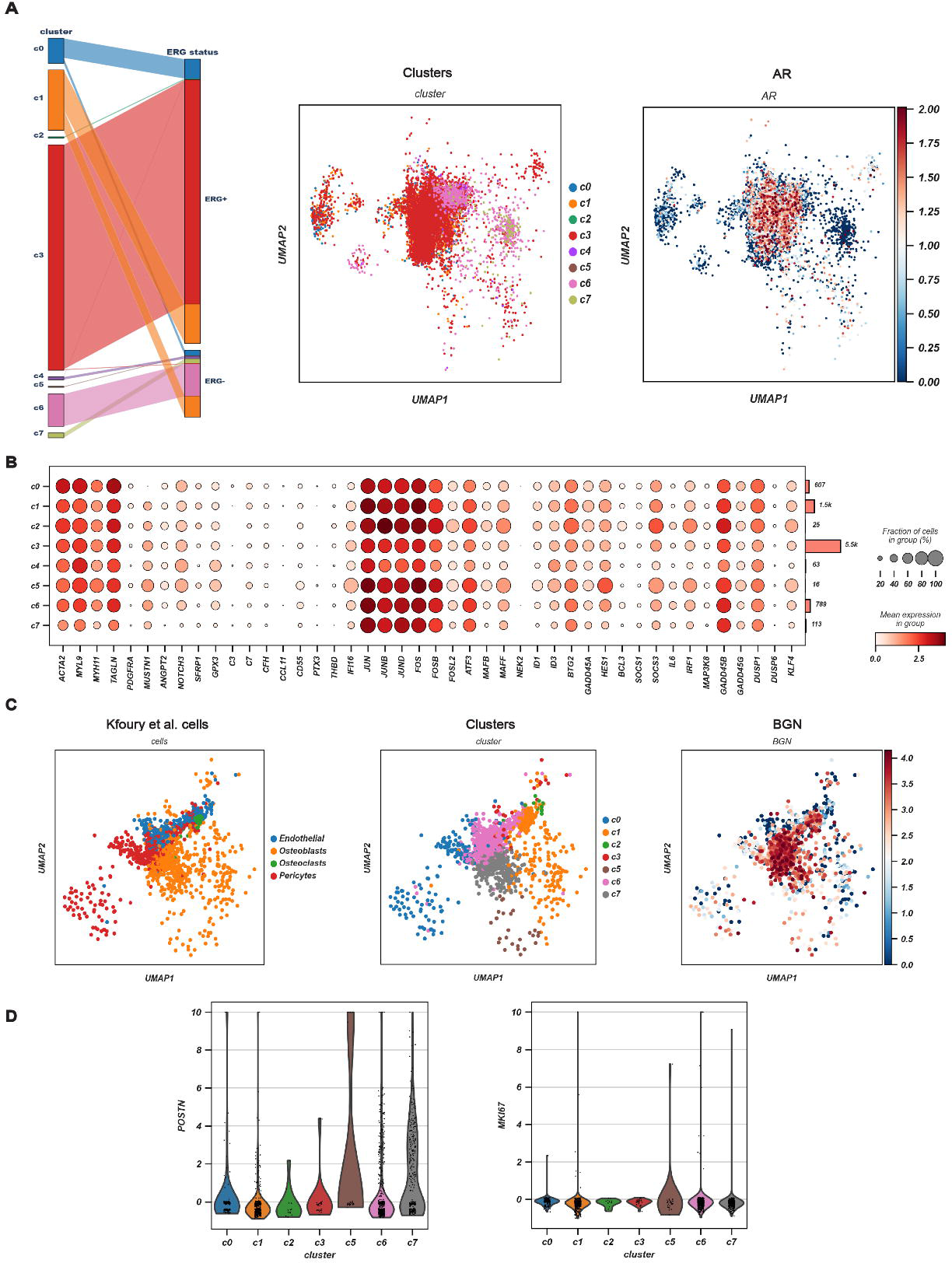
Analysis of human scRNA-seq data suggests the relevance of prostate mesenchyme in human PCa pathobiology. (A) Parallel categories plot showing the relationship between the mesenchymal clusters and ERG status (left). UMAP projection of the 8 mesenchymal clusters in the human scRNA-seq data (center) and *AR* expression in the human mesenchymal clusters (right). (B) Dot plot showing the mean expression of marker genes for common clusters c0-c2 in the human scRNA-seq data. (C) UMAP of the selected cell types from the bone metastasis scRNA-seq data derived from Kfoury et al., 2021 (left) and their associated annotation using the 8-mesenchymal cluster definition (middle). The expression of *BGN* across the 8 mesenchymal clusters is shown on the right. (D) Violin plots showing the mean expression of *POSTN* and proliferative marker *MKI67* across the mesenchymal clusters in the *Kfoury et al*. scRNA-seq cohort.

### Transcriptional similarities between the stroma of primary tumor and that of bone metastases

Analyses using scRNA-seq profiles of human PCa bone metastasis revealed transcriptional patterns similar to those present in the mesenchymal clusters from the *PRN* model (c5-c7), comprising more than 60% of bone stromal cells (**Figure 6C-D**) and showed high expression of *POSTN* and *MKI67* (**Figure 6D**), together with genes active in the bone microenvironment, such as *BGN* (**Figure 6C-D**). While the NEPC-related clusters were the most predominant in the metastatic microenvironment in bone, cells from c0 and c1 were also common, representing 9% and 27% of the total cells, respectively. Akin to primary cases, c6 and c7 in bone metastases expressed components of adaptive immune responses such like *C1Q A, B* and *C* (**Figure S5C**).

Taken together, these findings show functional and transcriptional similarities between the stroma of advanced PCa models and that of the bone microenvironment.

## Discussion

While different mutations in epithelial tumor cells partially explain the phenotypic and clinical heterogeneity of PCa, roughly one fourth of prostate tumors are genomically “quiet”^32^, indicating that additional undiscovered are key determinants of the biological behavior of PCa. Mesenchymal cells, which represent the predominant component of the microenvironment, have been suggested for decades to play a major role in this regard^5, 34, 35^. Recently, studies by *Karthaus et al.* and *Crowley et al.* described a detailed cluster analysis of mesenchymal cells in the mouse prostate by scRNA-seq, revealing a level of complexity greater than that suggested previously^22, 24^. Here, we analyzed in detail by scRNA-seq all mesenchymal cells utilizing all prostate lobes in the mouse prostate from several established GEMMs and corresponding WT mice. The significant and progressive increase in the mesenchymal cell component in increasingly aggressive GEMM models suggests a pivotal role of the stroma in tumor progression. We identified eight distinct stromal cell states that were defined by different gene expression programs and by underlying regulatory transcription factors. Three clusters represent fibroblasts states that are common to all genotypes, and they display conserved functional programs across all stages of tumor growth. On the other hand, we described five novel stromal cell states that are specifically linked to defined epithelial mutations and disease stages, setting the stage for a better understanding of mutation-specific epithelial-stromal interactions.

There is growing evidence that innate immunity and inflammation play a role in prostate and other cancers^35, 36, 37^. While the focus of this study was not on immune cells, we found a cluster of mesenchymal cells conserved across all genotypes in prostate mesenchyme expressing genes associated with immunoregulatory and inflammatory pathways and driven by transcription factors such as Nfkβ. Immune cells including tissue-resident macrophages are recruited and subsequently activated to modulate prostate tumorigenesis. In addition, stromal cells produce cytokines, chemokines and components of complement protein pathways^38, 39^. The complement system is an established component of innate immunity. Components of complement activation via the C3 alternative pathway were previously found to be activated by *KLK3* (a.k.a. PSA), with a special affinity for iC3b that in turn stimulates inflammation^40^. In addition, a pronounced expression of *Cd55* in common clusters, inhibits complement *C3* lysis^41^. The role of the complement as mediator of the stromal-immune crosstalk in c1 was also confirmed by the ligand-receptor analysis which showed significant interactions between *C3* and both ITGAM_ITGB2 and ITGAX_ITGB2 receptors in dendritic cells. This suggests that the expansion of cells expressing *C3* can stimulate innate immune response in the TME. Complex and bidirectional interactions between stroma and immune cells, mostly involve dendritic cells, monocyte/macrophages, and Tregs. Model-specific variations in the composition of the tumor immune microenvironment were seen, e.g. a prominent infiltration of monocytes/macrophages in the NP models. Further functional analyses of those interactions will reveal how the stroma influences the response to immunotherapy in PCa^42, 43, 44^.

Roughly half of prostate tumors have ETS translocations with *TMPRSS2* as the most frequent fusion partner^32^, one of the earliest alterations in prostatic carcinogenesis^45, 46, 47^. Yet, genetically engineered mouse models driven by the TMPRSS2-ERG fusion have little to no phenotype in the epithelium. Here, we found that induction of mesenchymal cell expansion is a significant early event in this model. We harmonized the eight murine clusters with human PCa cases sequenced using the same scRNA-seq approach. Strikingly, the mesenchyme associated with the TMPRSS-ERG translocation was conserved between mouse and human. Thus, epithelial ERG fusion in the mouse triggers early changes in the adjacent stroma, creating a TME that supports ERG-positive epithelial cells. Given the conservation of these mesenchymal clusters in humans, these findings show the role of this prevalent alteration in the pathogenesis of prostate cancer. It will be important to determine the prevalence of these stromal cluster associated with TMPRSS-ERG in patients of African descent, where the prevalence of this translocation is low^48^.

Stromal populations contribute to the structural and functional TME ecosystem through different autocrine and paracrine mechanisms. Among them, the stromal *AR* signaling cascade is known to influence prostate epithelial cells’ behavior at different stages of development and carcinogenesis^34, 49^. Stromal *AR* signaling may prevent invasion by maintaining an non-permissive TME for cell migration^50^. Indeed, loss of stromal AR was associated with upregulation of ECM-remodeling metalloproteinases (e.g., *MMP1*) and of *CCL2* and *CXCL8* cytokines, factors that promote invasion ^50, 51^. In the transgenic *Hi-MYC* and the testosterone+estradiol hormonal carcinogenesis models, stromal AR deletion, especially in smooth muscle cells favors prostate carcinogenesis^52^. In line with these observations, we show decreased mesenchymal *Ar* expression in the *PRN* model, which recapitulates late-stage PCa and progression towards neuroendocrine differentiation. Stromal *AR* may play a master role in committing and maintaining epithelial prostate cell identity in at least two ways. During development, its expression induces epithelial cells to differentiate into prostate cells, and during PCa development it prevents progression towards undifferentiated/neuroendocrine status.

Low expression of *Ar* in the *PRN* model was inversely associated with an increased expression of periostin (Postn), and *in situ* analyses confirmed that Postn-positive cells were enriched in areas of neuroendocrine differentiation. Stromal expression of periostin in PCa has been associated with decreased overall survival^53^ and higher Gleason score^54^. We show that stromal cells expressing *Postn* confer invasive ability to poorly differentiated/NE carcinoma. The increased expression of *Postn* and of genes typical for the bone microenvironment (e.g., *Bgn*) suggest that invasive PCa cells and the associated, invasion-primed mesenchyme modify the prostate TME to resemble that of bone, a common site of metastases in this malignancy. In this fashion, the primary site TME may pre-condition tumor cells for skeletal metastatic seeding. Importantly, we discovered shared characteristics between the stroma of the advanced/neuroendocrine GEMM and that of scRNA-seq stromal profiles from human bone metastases. Specifically, the bone stroma had a high frequency of *POSTN*+ cells,. It is yet to be determined whether these cells were inherently present in the bone microenvironment or expanded as a result of the metastatic process. These cells were also characterized by the expression of genes involved in osteoblast differentiation and proliferation like *RUNX2, BMP2, IGF1*, and *IGFBP3*^55^ together with Cadherin 11 (*CDH11*) previously found to induce PCa invasiveness and bone metastasis^56, 57^.

The role of complement is important not only in both modulating innate immunity but also invasion. A pronounced expression of *C1QA*, *B* and *C* was identified especially in models of advanced disease. *C1q* has been shown to promote trophoblast invasion^58^ as well as angiogenesis in wound healing^59^. This was in line with our previously published stromal signature derived from laser capture-microdissected (LCM) mesenchyme adjacent to high grade tumors that predicted lethality in an independent PCa cohort^8^. Three of the 24 signature genes were in fact *C1Q A*, *B,* and *C* suggesting that complement activation by the stroma plays a role in the invasive potential of aggressive prostate tumors (with diverse epithelial genetic alterations). The unexpected resemblance between PCa mesenchyme of locally aggressive tumors and that of bone metastases suggests that locally advanced PCa tumors prone to metastasize display a bone-like microenvironment. Since transcriptional profiles of stromal cells in aggressive models were conserved in the stroma of human localized high-grade tumors^8^, as well as in stroma of bone metastases in patients’ biopsies^60^, a broad set of cellular and molecular changes in the stromal cells may be either permissive or directly affect progression and metastatic disease.

Importantly, while our findings offer novel insights about the role of the stroma in mediating PCa progression and invasiveness, they also show a strong translational relevance. For instance, we have used the scRNA-seq transcriptional profiles of the PRN-derived mesenchymal clusters (c5-c7) to develop a robust and interpretable gene signature for predicting PCa metastases in a large cohort of patient samples with bulk transcriptomic profiles. This signature was also associated with worse progression-free survival in a separate cohort (TCGA) before and after adjusting for Gleason grade.

In summary, here we provide a molecular compendium of mesenchymal changes during PCa progression in genetically engineered mice that generalize to humans. Specifically, in the early phases of prostate carcinogenesis, we provide evidence that the TMPRSS-ERG translocation reprograms the mesenchyme which in turn may sustain progression. Moreover, in advanced PCa models we found transcriptional mesenchymal programs linked to metastasis, some of them in common with the bone microenvironment to which PCa cells metastasize. The findings from those murine models have been validated and confirmed using publicly available and internally generated scRNA-seq data from ERG+ human tumors and PCa bone metastases. Collectively these data in mice and human provide unambiguous evidence for marked shifts in stromal composition accompanying PCa progression and driven in a genotype-specific manner, ultimately implicating mesenchymal changes as major contributors of both PCa progression and phenotypic diversity to a degree not previously appreciated.

## Methods

### Genetically engineered mouse models of prostate cancer

We focused on three models of prostate cancer that reflect the most common mutations in human localized disease, plus a fourth model that recapitulates the transition to NEPC. The choice of these models was also taken to reflect different stages of the disease.

Specifically, the TMPRSS2-ERG (*T-ERG)* fusion model has an N terminus-truncated human ERG together with an ires-GFP cassette into exon 2 of the mouse Tmprss2 locus^11^, displays a minimal epithelial phenotype in the mouse, and was chosen since it represents the most frequent mutation in human prostate cancers^32^. Pten knock-out (*NP*) mice develop high grade PIN with areas of invasion. To obtain Nkx3.1creERT2;Pten^f/f;^ *EYFP^f/f^* (*NP*^12^, *Nkx3.1^creERT2^* driver was crossed to the conditional allele for *Pten* (Pten^flox/flox^) with loxP^61, 62^. For induction of Cre activity in NP mice, tamoxifen (Sigma Cat #T5648) (or corn oil alone) was delivered by IP injection (225mg/kg) for 4 consecutive days, to mice at 2 months of age. Six months later NP mice were sacrificed and analyzed. *Hi-MYC*^14^ shows both PIN and microinvasion. PRN mice carry the *MYCN* transgene, *Pten* and *Rb1* homozygous floxed^63^. Additionally, non littermate WT mouse strains, *FVB/N* (Charles River, CRL), *C57BL/6* and *B6129SF2/J* Laboratories-JAX) were used.

All animals used in this study received humane care in compliance with the principles stated in the Guide for the Care and Use of Laboratory Animals (National Research Council, 2011 edition), and the protocol was approved by the Institutional Animal Care and Use Committee of Weill Cornell Medicine, Dana-Farber Cancer Institute and Columbia University Irving Medical Center.

The number of GEMMs and their WTs and/or littermates are provided in **Table S1**.

### Description of human prostate cancer specimens

Human prostate tissue specimens were obtained from patients undergoing radical prostatectomy at Weill Cornell Medicine under Institutional Review Board approval (WCM IRB #1008011210). These included 9 samples (3 ERG-negative and 6 ERG-positive cases). The clinical and molecular characteristics of these patients are provided in **Table S5**. Immediately after surgical removal, the prostate was sectioned transversely through the apex, mid, and base^64^. Tissue for scRNA-seq was placed in RPMI medium with 5% fetal bovine serum (FBS) on ice, and quickly transported for single-cell RNA sequencing. A small portion of the regions of interest, including the areas selected for single-cell RNA sequencing, index lesion, and contralateral benign peripheral zone, was concomitantly frozen in optimal cutting temperature (OCT) compound, cryosectioned, and a rapid review was performed by a board-certified surgical pathologist (BR) to provide a preliminary assessment on the presence of tumor, normal epithelium, stroma near and away from the tumor. Adjacent tissue was processed by formalin fixation and paraffin embedding, followed by sectioning, histological review, histochemistry (trichrome stain), and immunostaining^65^.

### Isolation of single cells for RNA-Seq

Dissociated murine prostate cells were prepared as described previously^66^. Briefly, mouse prostate tissues were digested in Advanced DMEM/F12/Collagenase II (1.5mg/ml)/Hyaluronidase VIII (1000 u/ml) (Thermo Fisher Scientific) plus 10 μM Y-27632 (Tocris) for 1 hour at 37°C with 1500 rpm mixing, continuously agitated. Subsequently, after centrifuging at 150 g for 5 min at 4°C, digested cells were suspended in 1 ml TrypLE with 10 µM Y-27632 and digested for 15 min at 37°C and neutralized in aDMEM/F12/FBS (0.05%). Dissociated cells were subsequently passed through 70 μm and 40 μm cell strainers (BD Biosciences, San Jose, CA) to obtain a single cells suspension. Samples were resuspended in 1x PBS and sorted by Flow Cytometry (Becton-Dickinson Aria II and/or Becton-Dickinson Influx) for 4′,6-diamidino-2-phenylindole (DAPI) to enrich for living cells.

Similarly, human prostate tissues were first digested in aDMEM/F12/Collagenase II (1.5mg/ml)/Hyaluronidase VIII (1000 u/ml; Thermo Fisher Scientific) plus 10 μM Y-27632 (Tocris) for 1 hour at 37°C with 1500 rpm mixing, continuously agitated. Subsequently, after centrifuging at 150 g for 5 min at 4°C, digested cells were suspended in 1 ml TrypLE with 10 µM Y-27632 and digested for 15 min at 37°C and neutralized in aDMEM/F12/FBS (0.05%). Dissociated cells were subsequently passed through 70 μm and 40 μm cell strainers (BD Biosciences, San Jose, CA) to get single cells. Samples were resuspended in 1x PBS and sorted for DAPI to enrich living cells.

Barcoded cDNA libraries were created from single-cell suspensions using the Chromium Single Cell 3’ Library and Gel Bead Kit, and Chip Kit from 10x Genomics^67^, according to manufacturer recommendations. Briefly, depending on the GEMMs and human samples used in this study, 8,000-16,000 cells were targeted for 3’ RNA library preparation, multiplexed in an Illumina NovaSeq 6000, and sequenced at an average depth of 25,000 reads per cell.

### Quantification and preprocessing of single-cell RNA sequencing data

Expression matrices were generated from raw Illumina sequencing output using CellRanger. Bcl files were demultiplexed by bcl2fastq, then reads were aligned using STAR^68^. All data collected from mouse models were aligned to GRCm38 reference transcriptome. To identify cells with trans-gene expression, we indexed and aligned to human ERG and GFP from the *T-ERG* model, human MYC from the *Hi-MYC* model, and human MYCN from the *PRN* model. Human data were aligned to GRCh38. Alignment quality control was performed using the default CellRanger settings. Expression matrices from the different mouse models were converted to AnnData objects and concatenated into a single count matrix using the Scanpy library (version 1.5) in Python (version 3.8)^33^. Similarly, the expression matrices from the nine human samples were concatenated into a single count matrix. The raw mouse and human scRNA-seq count matrices were preprocessed as follows: cells with low UMIs (unique molecular identifiers) count (<400) and low number of expressed genes (<300) were removed. Subsequently, genes that were expressed in three or fewer cells and cells containing more than 20% mitochondrial transcripts were removed after visualizing the distribution of fraction of counts from mitochondrial genes per barcode^69^. Contributions from total count, mitochondrial count, and cell cycle were corrected by linear regression. The resulting matrix was then log1p transformed^69^. Finally, the top 4,000 genes were selected by coefficient of variation according to the method described in^67^, and genes were scaled to mean zero and unit variance^69^.

### Embedding of scRNA-seq expression matrix by deep generative modeling

We computed batch-corrected embeddings as follows. We fit our data using a conditional variational autoencoder^70^. Specifically, we used the negative binomial counts model included in the single-cell variational inference (scVI) Python package^71^. We model (a) a nuisance variable that represents differences in capture efficiency and sequencing depth and serves as a cell-specific scaling factor, and (b) an intermediate value that provides batch-corrected normalized estimates of the percentage of transcripts in each cell that originate from each gene. Our model is implemented in Python using the PyTorch library (v1.7.0)^72^ and was run on a NVIDIA RTX A4000 GPU.

### Clustering and data visualization

A nearest neighbor graph was constructed with Euclidean metric from the batch-corrected scVI embeddings, then cells were partitioned by the Leiden clustering algorithm^73, 74^. Partition-Based Graph Abstraction (PAGA) was computed from the Leiden partition^75^ and was used to initialize the Uniform Manifold Approximation and Projection (UMAP) algorithm which projected the data into 2D space^76^.

### Identification of stromal cells

For both the mouse (101,853 cells) and human (83,080 cells) scRNA-seq datasets, we excluded cells of lymphoid, endothelial, and neural origin based on Leiden clustering at resolution 1.0 and the expression of associated lineage markers (**Table S2**). Subsequently, we identified stromal cells as cells expressing either of two canonical mesenchymal marker gene sets: ‘mesenchymal 1’ or ‘mesenchymal 2’, respectively (**Table S2**)^22, 24, 77, 78, 79^. The resulting mesenchymal datasets for the mouse and human scRNA-seq data included 8,574 and 8,628 cells, respectively. These mesenchymal cells were then clustered using the Leiden algorithm to identify different mesenchymal subclusters. Specifically, at resolution 0.05, the Leiden clustering reflected the separation of *Mesenchymal* and *Smooth Muscle Cells/Myofibroblasts* subtypes. We increased resolution in increments of 0.05, inspecting the biological plausibility of new clusters until resolution 0.6 (**Table S3**), after which higher resolution produced new clusters with differences dominated by noise^73^.

### Identification and annotation of immune cell types

In the mouse scRNA-seq data, cells from the immune compartment (42,431 cells) were also clustered using the Leiden algorithm. The resulting clusters were then annotated to different immune cell types based on the expression of known markers genes. These included B cells (expressing *Cd79a, Cd79b, Cd74, Cd19,* and *Cd22*), CD4+ T lymphocytes (expressing *Cd4, Cd2, Cd28,* and *Trac*), NK or cytotoxic T cells (expressing *Xcl1, Nkg7, Gzmb, Klrc1,* and *Klrc2*), Tregs (expressing *Foxp3, Ctla4, Tnfrsf4*, and *Tnfrsf18*), dendritic cells (expressing *Ccl17, Ccr7, Xcr1*, and *Cd207*), and monocytes or macrophages (expressing *Cd68, Cd74, Cxcl2,* and *Lgals3*).

### Differential expression testing

For differential expression (DE) testing, we used a two-part generalized linear model (hurdle model), MAST, that parameterizes stochastic dropout and the characteristic bimodal distribution of single cell transcriptomic data^20^. DE was performed by comparing cells from each cluster to pooled cells from all other clusters (**Table S3**).

### Gene regulatory network inference (GRN)

Gene regulatory network activity was inferred from the raw counts matrix by pySCENIC (v0.10.3) ^19^. Specifically, coexpression modules between transcription factors (TFs) and their candidate targets (regulons) were inferred using the Arboreto package (GRNBoost2) and pruned for motif enrichment to separate indirect from direct targets^19, 80^. The activity of each regulon in each cell was then scored using the Area Under the ROC curve (AUC) calculated by the *AUCell* module from pySCENIC package^19, 80^. Cluster-specific regulons were identified as those with AUCell Z-score >1 for each mesenchymal cluster.

### Ligand-receptor analysis

We performed ligand-receptor (LR) interaction analysis using CellChatDB and CellChat R tool (version 1.1.3) to predict cell-cell interactions within the tumor microenvironment^81^. Cell communication networks were inferred by first identifying differentially expressed ligands and receptors between the different mesenchymal clusters, immune cell types, and the epithelium. The probabilities of these interactions on the ligand-receptor level were computed using the default ‘trimean’ method setting the average expression of a signaling gene to zero if it is expressed in less than 25% of the cells in one group. Notably, we corrected for the effect of cluster size (number of cells) when calculating the interaction probabilities. Additionally, we summarized the ligand-receptor interaction probabilities within each signaling pathway to compute pathway-level communication probabilities. Cell-cell communication networks were then aggregated by summing the number of interactions or by averaging the previously calculated communication probabilities. To compare the signaling patterns between mutants and wild types (**Figure 1E**), we first performed differential expression analysis between all the mutants versus wild types in each of the three compartments (stroma, epithelium, and immune). Upregulated LR pairs were identified if each had a log fold change (logFC) above 0.1 in the senders and receivers, respectively. Finally, we extracted the mutant-specific LR pairs as those with up-regulated ligands and receptors in the mutants compared to wild types and vice versa. In this analysis, we used a p-value threshold of 0.01 to determine significant interactions.

### Label transfer from mouse to human scRNA-seq data

To transfer the stromal cluster labels from the mouse to human data, human gene symbols were first converted to their mouse counterparts then both datasets were restricted to overlapping genes. Label transfer was performed using *‘ingest’*^33^ which maps the labels and embeddings fitted on an annotated reference dataset to the target one. Specifically, we used the scRNA-seq data from the mouse *T-ERG* model as reference for the human ERG-positive cases and those from the remaining mouse models as reference for the human ERG-negative cases. Finally, we computed the ranking of differentially expressed genes in each cluster versus the remaining ones using t-test.

### Processing of human bone metastases scRNA-seq data

The raw count matrix of the scRNA-seq dataset previously reported by Kfoury et al.^60^ was retrieved from the Gene Expression Omnibus (GEO). This dataset included 25 bone metastasis samples derived from PCa patients, of which 9 samples were derived from solid metastasis tissue. Further analysis was limited to these 9 samples (16,993 cells). The data was preprocessed by keeping cells with at least 200 expressed genes and less than 15% mitochondrial transcripts (16,536 cells). Subsequently, cells were normalized by the total counts over all genes followed by log scaling and regressing over the total counts per cell and percentage of mitochondrial genes to reduce unwanted variation. The top 4000 highly variable genes were selected and the resulting matrix was then scaled to unit variance and zero mean. Since this particular analysis was intended to explore the transcriptional and functional similarities between the primary tumor stroma and the stroma of bone metastasis, we further limited the analysis to the cells previously annotated by the authors as osteoblasts, osteoclasts, endothelial cells, and pericytes (1,872 total cells). Finally, the embeddings and stromal cluster labels were projected onto this dataset using the mouse stroma scRNA-seq dataset as reference and following the same steps mentioned above.

### Development of the PRN signature to predict metastasis in prostate cancer patients

We collected and curate gene expression profiles from different datasets comprising 1239 primary tumor samples from PCa patients with information about metastastatic events. These datasets included six publicly available datasets (GSE116918, GSE55935, GSE51066, GSE46691, GSE41408, and GSE70769, together with a seventh dataset available from Johns Hopkins University, referred to as the natural history cohort^82^. The expression profiles from each dataset were normalized, log2-scaled, then z-score transformed (by gene) separetly. Subsequently, we mapped probe IDs to theircorresponding gene symbols and kept only the genes in common between all datasets (12761 genes).

The 1239 samples were joined together then splitted into 75% training (n=930) and 25% testing (n=309) using a stratified sampling approach to ensure an equal representation of important variables including the original datasets, Gleason grade, age, tumor stage, and prostate-specific antigen (PSA) levels. Quantile normalization was applied to both the training and testing sets separately. The training set was used for training a classifier that can predict metastasis using the k-top scoring pairs (k-TSPs) algorithm, which is a rank-based method whose predictions depend entirely on the ranking of gene pairs in each sample^83, 84^. Based on the average log fold change (logFC), we divided the markers of the PRN clusters into positive (average logFC > 0) and negative (average logFC <0) markers. We then paired the top positive and negative markers (100 genes each) together to build a biological mechanism representing the PRN mesenchyme (30000 pairs). Each pair consists of two genes, one is up- and another is down-regulated in the PRN mesenchyme. This mechanism was then used as a priori biological constraint during the training of the k-TSPs algorithm^85^, and the resulting signature was evaluated on the indepedent testing set.

Additionally, we evaluated the prognostic relevance of this signature in the TCGA cohort which included 493 primary PCa samples. First, we built a logistic regression model using the 26 genes comprising the PRN signature and used this model to generate a probability score for progression-free survival (PFS) in each patient. We then binarized these probabilities into predicted classes and compared their PFS probability using Kaplan-Meier survival analysis^86^. Finally, we calculated the hazard ratio of the signature prediction probability scores after adjusting for Gleason grade using a multivariate Cox proportional hazards model^87^.

### Histopathology studies

Following radical prostatectomy, human prostates were submitted for gross pathological assessment and sectioning, with ischemic time less than 1 hour. The prostate specimen was serially sectioned from apex to base into 3-5 mm slices. In prostates with grossly identifiable tumor, a 5 mm biopsy punch was taken from the area of tumor, an area adjacent to the tumor, and an area distant (>2 slices away) from the tumor. In prostatectomy specimens where tumor was not definitively grossly visible, these areas were approximated by anatomic correlation of the MRI findings and targeted biopsies with the highest tumor grade (as described in ^64^).

The prostate slices were fixed in 10% buffered formalin, embedded in paraffin blocks, and hematoxylin & eosin (H&E)-stained slides were created, per routine clinical pathologic assessment.^51,52^. Upon evaluation of the H&E slides, a urologic pathologist (BDR) confirmed that the punched area of tumor, area adjacent to tumor, and area distant to tumor were accurately represented based on the histology of the areas surrounding the punched area. Prostate from WT and GEMM mice were dissected. One half of the prostate from GEMMs was utilized for scRNA-seq (see above). The contralateral half was fixed in 10% buffered formalin and embedded in paraffin blocks, sections were cut, and hematoxylin & eosin (H&E)-stained slides^88, 89^. Collagen deposition in the different GEMMs was assessed by Masson’s trichome staining^90, 91^, followed by collagen deposition quantification digitally performed using HALO (Indica Labs, v3.3.2541, Albuquerque, US). A HALO-based digital classifier was developed to identify collagen, epithelium, muscle fiber, and background regions on the digital images. Percentages of collagen deposition were then quantified and compared using unpaired t-test. Immunohistochemical stainings were used to confirm the expression of the GEMMs proteins.

Immunohistochemistry to interrogate for panel markers (**Table S6**) was performed on 5-μm-thick formalin-fixed paraffin-embedded tissue (FFPE) of (i) human PCa and (ii) GEMMs sections using previously-established protocols^88, 92, 93^.

Multiplexed immunohistochemistry (mIHC) was performed by staining 5-μm-thick FFPE core biopsy sections in a BondRX automated stainer, using published protocols^94, 95, 96^. One panel of primary antibody/fluorophore pairs was applied to all cases along with Antibody/Akoya Opal Polaris 7-Color Automated IHC Detection Kit (NEL871001KT), and Opal Polymer Anti-Rabbit HRP kit for secondary antibody (ARR1001KT) fluor combinations were utilized as follows (**Table S6**). The order of processing slides was as follow: primary antibody incubated for 30 minutes; Blocking for 5 minutes with Akoya Blocking/Ab Diluent; Opal Polymer Anti-Rabbit HRP incubated for 30 minutes; Opal 480-690 incubated for 10 minutes; Leica Bond ER1 solution incubated for 20 minutes. All slides were also stained with DAPI for nuclear identification.

### Acquisition and Computational Analysis of Multiplexed Immunofluorescence Images

Whole slide images of hematoxylin and eosin, trichrome and mIHC sections were acquired using the Vectra Polaris Automated Quantitative Pathology Imaging System (Akoya Biosciences, Hopkinton, MA)^97^. Images were processed by linear spectral unmixing and deconvolved^98^. Cells were segmented and a human-in-the-loop HALO random forest classifier was trained with labels from a pathologist to select stromal cells. Subsequently, these stromal regions of the entire prostate surrounding glands in WT and GEMMs mice were preprocessed and analyzed using PathML (https://github.com/Dana-Farber-AIOS/pathml)99 to generate a single cell counts matrix containing statistics summarizing the expression of each protein in each cell together with the cell size, coordinates, and eccentricity. To address technical artifacts in the segmentation results, DAPI-negative cells were filtered out. A nearest-neighbor graph was constructed from the counts matrix using Euclidean metric as implemented in the Scanpy package^33^. This graph was clustered using the Leiden algorithm^73^ to identify subpopulations of cells and low-quality cells. Cells were projected to two dimensions and visualized using the UMAP algorithm^76^. A binary label indicating the presence/absence of each protein was created by thresholding markers for positive or negative signal with pathologist assistance.

### RNA Extraction and bulk RNA-seq analysis of murine PRN tumors

Fresh-frozen prostate tumors from *PRN* mice were cryopreserved in OCT. Histological evaluations and quantifications (including H&E staining and IHC staining) were performed by a board-certified, genitourinary pathologist (BDR) who was blinded to animal genotypes and followed criteria that have previously been described^100^. Regions of the OCT block identified by a blinded pathologist as NEPC or adenocarcinoma were cored using a biopsy punch. RNA extraction was performed on the frozen cores using the Maxwell 16 LEV simplyRNA Tissue Kit (Promega, AS1280). Specimens were prepared for RNA-seq as described above and RNA quality was verified using Agilent Bioanalyzer 2100 (Agilent Technologies). Paired-end, 50 × 2 cycles sequencing was performed on the HiSeq 4000 instrument. Quality control of raw sequencing reads was performed using FastQC (Babraham Bioinformatics). Low-quality reads were removed using Trimmomatic^101^ with a sliding window size of 4 bp and a quality threshold of 20. The resulting reads were aligned to mm10 using STAR^68^. Reads were sorted and indexed using SAMtools^102^. Transcript abundance was calculated using HTSeq^103^. Differential gene expression was assessed using DESeq2^104^.

### Generation of murine Normal Associated Fibroblasts (NAFs)

Prostate tissues derived from 3-month-old C57/BL6 male mice were minced in apron 1 mm pieces and placed in p100 using DMEN+5%FBS+5%NuSerum+1%Gln+1%P/S+10nMDHT. The fibroblasts were attached to the plate within 48-96 hours and the chunks were removed. Then the immortalization was performed using Retrovirus with zeocin resistance and expression of SV40 T antigen (pBabe-Zeo-LT-ST). NAFs were cultured in normal DMEM+10%FBS+1%Gln+1%P/S.

### RNA knockdown

For lentiviral shRNA transduction, mouse NAFs were transduced using lentiviruses containing shRNA constructs against *Postn* with 10 mg/ml polybrene (Sigma, TR-1003-G). shPostn1 (F primer: CACCGGGCCATTCACATATTCCGAGAACTCGAGTTCTCGGAATATGTGAATGGCTTTTTG; R primer: GATCCAAAAAGCCATTCACATATTCCGAGAACTCGAGTTCTCGGAATATGTGAATG GCCC) shPostn2 (F primer: CACCGGCCACATGGTTAATAAGAGAATCTCGAGATTCTCTTATTAACCATGTGGTTTTTG; R primer: GATCCAAAAACCACATGGTTAATAAGAGAATCTCGAGATTCTCTTATTAACCATGTG GCC).

### Migration (invasion) assay

For Boyden chamber assays, 100,000 NAFs infected with control of Periostin-directed shRNAs (shCtrl, shPostn1 or shPostn2) were seeded into a 24-well plate in culture media for 24 hours. Cells were washed twice for 15 minutes in minimal media (DMEM (Thermo Fisher, 31053036) with 1× penicillin/streptomycin (Gibco, 15140-122), 1× GlutaMAX (Gibco, 35050-061) and 10 mM HEPES (Gibco, 15630-130). Cell culture inserts (Millipore, #MCEP24H48) were coated with Matrigel (Corning, 354230) diluted 1:10 in PBS, and incubated for 2 hours +37°C. 22rv1 shRb1 NMYC cells were harvested, washed twice for 3 minutes in minimal media, and seeded in triplicate at a density of 75,000 cells/insert in 200ul. Inserts were placed into empty 24-well plates and incubated for 15 min at +37°C and 5% CO2 before transferring into the test conditions, minimal media was used as a negative control and minimal media supplemented with 10% charcoal-stripped serum (Gibco, A33821-01) was used as a positive control. Cells were then allowed to migrate at +37°C for 6 hours. Filters were fixed using 4% PFA/PBS, washed with PBS, and stained using Hoechst before washing, cleaning and mounting using Fluoromount-G (SouthernBiotech, 0100-01). Cells that had migrated through the filter were quantified (5 fields of view per filter) and normalized to the negative control.

## Data and code availability

Single-cell RNA-seq data have been deposited at Zenodo for the purpose of peer review. Data will be deposited at GEO and made publicly available as of the date of publication. Microscopy data reported in this paper will be shared by the lead contact upon request. All original code has been deposited at GitHub and is publicly available using this link: https://github.com/MohamedOmar2020/pca_TME. Any additional information required to reanalyze the data reported in this paper is available from the lead contact upon request.

## Acknowledgements

We are grateful to the following facilities and individuals at Weill Cornell Medicine for their contributions to this work: Research Animal Resource Center, the Genomics Cores (Jenny Xiang, Dong Xu, Chendong Pan, Aihong Liu), the Translational Research Program in the Department of Pathology and Laboratory Medicine (Bing He, Yifang Liu, Leticia Dizon), Flow Cytometry Core Facility (Jason McCormick), Clinical Genomics Lab (James Solomon). This work was supported in part by the National Cancer Institute (Prostate Cancer SPORE Grant P50CA211024 [M.L., H.P., R.C., M.O., S.R., C.R.], P01 CA265768-01 [M.L., H.P., R.C., M.O., S.R., C.R.], R01CA200859 [L.M.], R01CA173481 [C.A.S], R01CA183929 [C.A.S], T32CA260293 [M.K.A]), and by an AIRC Fellowship for Abroad “Ezio, Maria e Bianca Panciera” [F.P.].

## Author information

These authors contributed equally: Hubert Pakula, Mohamed Omar, and Ryan Carelli.

**Department of Pathology and Laboratory Medicine, Weill Cornell Medicine, New York, NY 10021, USA**

Hubert Pakula, Mohamed Omar, Ryan Carelli, Filippo Pederzoli, Giuseppe Nicolò Fanelli, Tania Pannellini, Lucie Van Emmenis, Silvia Rodrigues, Caroline Fidalgo-Ribeiro, Pier V. Nuzzo, Nicholas J. Brady, Madhavi Jere, Caitlin Unkenholz, Mohammad K. Alexanderani, Francesca Khani, Matthew B. Greenblatt, David S. Rickman, Christopher E. Barbieri, Brian D. Robinson, Luigi Marchionni, Massimo Loda

**Sandra and Edward Meyer Cancer Center, Weill Cornell Medicine, Belfer Research Building, 413 East 69th Street, New York, NY 10021, USA**

Francesca Khani, Christopher E. Barbieri, Brian D. Robinson

**Department of Urology, Weill Cornell Medicine, New York, NY 10021, USA.**

Francesca Khani, Christopher E. Barbieri, Brian D. Robinson

**Department of Oncologic Pathology, Dana-Farber Cancer Institute and Harvard Medical School, 450 Brookline Ave, Boston, MA, 02215, USA.**

Massimo Loda

**Departments of Molecular Pharmacology and Therapeutics, Urology, Medicine, Pathology & Cell Biology and Systems Biology, Herbert Irving Comprehensive Cancer Center, Vagelos College of Physicians and Surgeons, Columbia University Irving Medical Center, New York, NY 10032, USA.**

Francisca Nunes de Almeida, Cory Abate-Shen

**Department of Laboratory Medicine, Pisa University Hospital, Division of Pathology, Department of Translational Research and New Technologies in Medicine and Surgery, University of Pisa, Pisa 56126, Italy:**

Giuseppe Nicolò Fanelli

## Contributions

H.P., R.C., M.O. and M.L. conceived and designed the study.

H.P., M.J., L.V.M, S.R., C.F-R., N.J.B., C.U., M.K.A., F.N-d-A. collected mouse and human data. C.E.B., T.P., F.K., M.K.A., B.R. provided study materials or patient samples and H.P., R.C., M.O., G.N.F., F.P. T.P., L.V.M, N.J.B, C.A.S, M.B.G., F.K., B.R., P.V.N., D.S.R., M.L., L.M., investigated, analyzed and interpret data, R.C., M.O., L.M., processed and analyzed single-cell data, T.P., G.N.F., C.U. performed immunohistochemistry. H.P., R.C., F.P., M.O., L.M., C.B., D.S.R., M.L wrote, reviewed manuscript. All authors edited and approved the manuscript.

## Corresponding author

Massimo Loda

## Ethics declarations

The authors declare no competing interests.

## Supplemental tables titles and legends

**Table S1. Genetically engineered mouse models and corresponding wild types used in the present study.**

Number of mice sequenced for each model and wildtype used in the present study, together with associated numbers of sequenced cells and transcripts.

**Table S2. Canonical gene markers used for cell type identification.**

List of canonical lineage gene markers used for cell type identification.

**Table S3. Differentially expressed genes (DEGs) according to designated clusters.**

List of differentially expressed genes for each of the newly identified mesenchymal clusters.

**Table S4. The PRN signature for predicting prostate cancer metastasis.**

The signature consists of 13 gene pairs with each including a gene up- and another down-regulated in the PRN mesenchyme (c5-c7). Pairs vote for metastasis if the 1^st^ gene is overexpressed relative to second. A patient with >=7 votes will be predicted to have metastasis.

**Table S5. Human PCa samples.**

List of human PCa samples used in the present study for scRNA-seq.

**Table S6. Multiplex immunohistochemistry antibody panels for cluster validation.**

## Supplemental figures titles and legends

**Figure S1. GEMMs have increased stromal formation compared to their WT counterparts.**

(A) Representative images of Masson’s trichrome show the increasing collagen deposition in tumor models according to the aggressiveness of the disease (left panel). HALO-based digital classifier categorizes tissue into collagen, epithelium, muscle fiber, and background components (middle panel). Collagen deposition is significantly enriched in *NP*, *Hi-MYC* and *PRN* models compared to their respective WTs (ns = not significant; * = p<0.05; ** = p<0.01; 2-tailed unpaired t-test; right panel).

(B) Representative H&E and IHC images showing stromal reaction in the presence of characteristic GEMM proteins.

Magnification for all images 200x. Scalebar: 300µm.

**Figure S2. Heatmap of the binarized regulon activity in the different stromal clusters in the mouse scRNA-seq data.**

The activity of each regulon in each cell was computed using the AUCell algorithm within the SCENIC workflow and then binarized using automatic cutoffs into active (black) or non-active (white). Shown is heatmap of the binarized activity of significant regulons (right) across all mesenchymal cells grouped by their corresponding cluster (top).

**Figure S3. Complex interactions between prostate cancer mesenchyme and different Immune cells.**

(A) Stacked bar plots showing the proportion of different immune cell types in the different mouse models.

Chord diagram showing the significant interactions between stromal and immune cells mediated by the CCL signaling pathway. Communication probsbilities were calculated after adjusting for the number of cells in each cluster.

(B) Chord diagrams of the significant signaling (ligand-receptor level) received by the stroma from Tregs (left) and monocytes/macrophages (right). The size of the inner bars represents the signal strength received by the stromal clusters. Communication probsbilities were calculated after adjusting for the number of cells in each cluster

**Figure S4. Signaling networks between the different mesenchymal clusters.**

(A) Significant ligand-receptor interactions across the 8 mesenchymal clusters.

(B) Significant stromal-stromal interactions mediated by the WNT signaling pathway.

(C) Significant stromal-stromal interactions mediated by the POSTN signaling pathway.

The size of the inner bars represents the signal strength received by their targets.

**Figure S5. C1Qs complement proteins activation in mesenchymal cells supports growth invasion and metastasis of poorly differentiated tumors in humans and PRN model.**

(A) Dot plot (left) and violon plots (right) showing the expression of *C1qa, C1qb and C1qc* across the stroma of mouse models. The highest expression of these complement genes is observed in the stroma of PRN (highlighted in the red bracket). Size of the dots represents the percentage of cells expressing the gene and color intensity represents the average expression level. Violin plots showing expression of *C1QA, C1QB* and *C1C* across the stromal clusters in the scRNA-seq data from primary PCa (B) and bone metastases data from the *Kfoury et al*. scRNA-seq cohort (C).

**Figure S6. Multispectral staining of prostate cancer tissue from the PRN mouse model showing AR and Periostin expression.**

(A) Expression of Periostin (upper) and AR (lower).

(B) AR+ and Periostin+ stromal cells (PanCK-) were identified by thresholding the intensities of PanCK, AR and Periostin.

(C) Heatmap of the spatial neighborhoods enrichment scores of AR+ and Periostin+ epethlial and stromal cells. Scores are based on proximity on the connectivity graph of cell clusters. The number of observed events is compared against 1000 permutations followed by z-score transformation.

