## Supplementary Figures for "Distinct mesenchymal cell states mediate prostate cancer progression"

### Figure S1

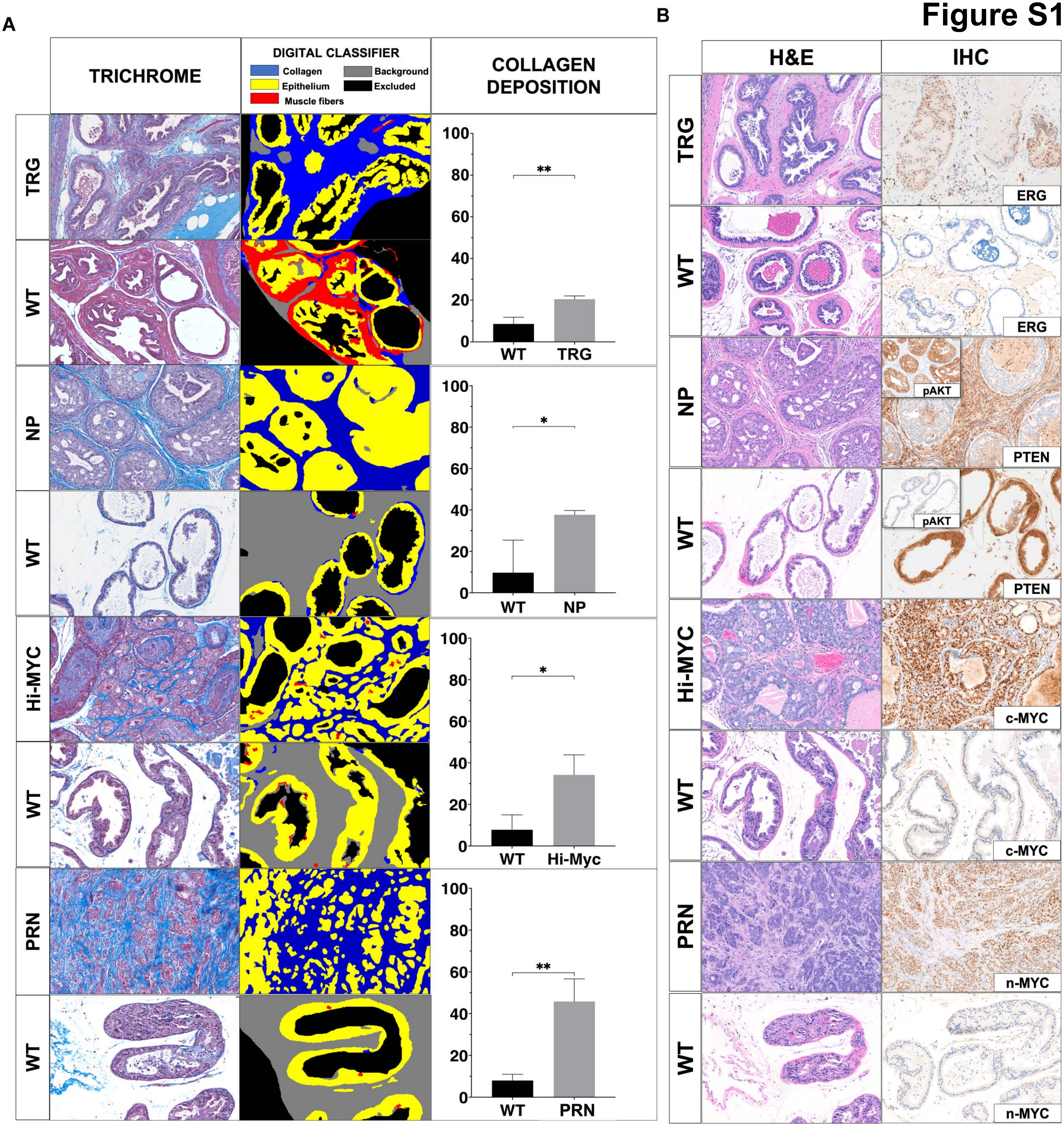
A
TRICHROME
DIGITAL CLASSIFIER
Collagen
Background
Epithelium
Excluded
Muscle fibers
COLLAGEN DEPOSITION
TRG
WT
NP
Hi-MYC
PRN
100
80
60
40
20
0
**
*
WT
TRG
NP
Hi-Myc
PRN
B
H&E
IHC
ERG
pAKT
PTEN
c-MYC
n-MYC

### Figure S2

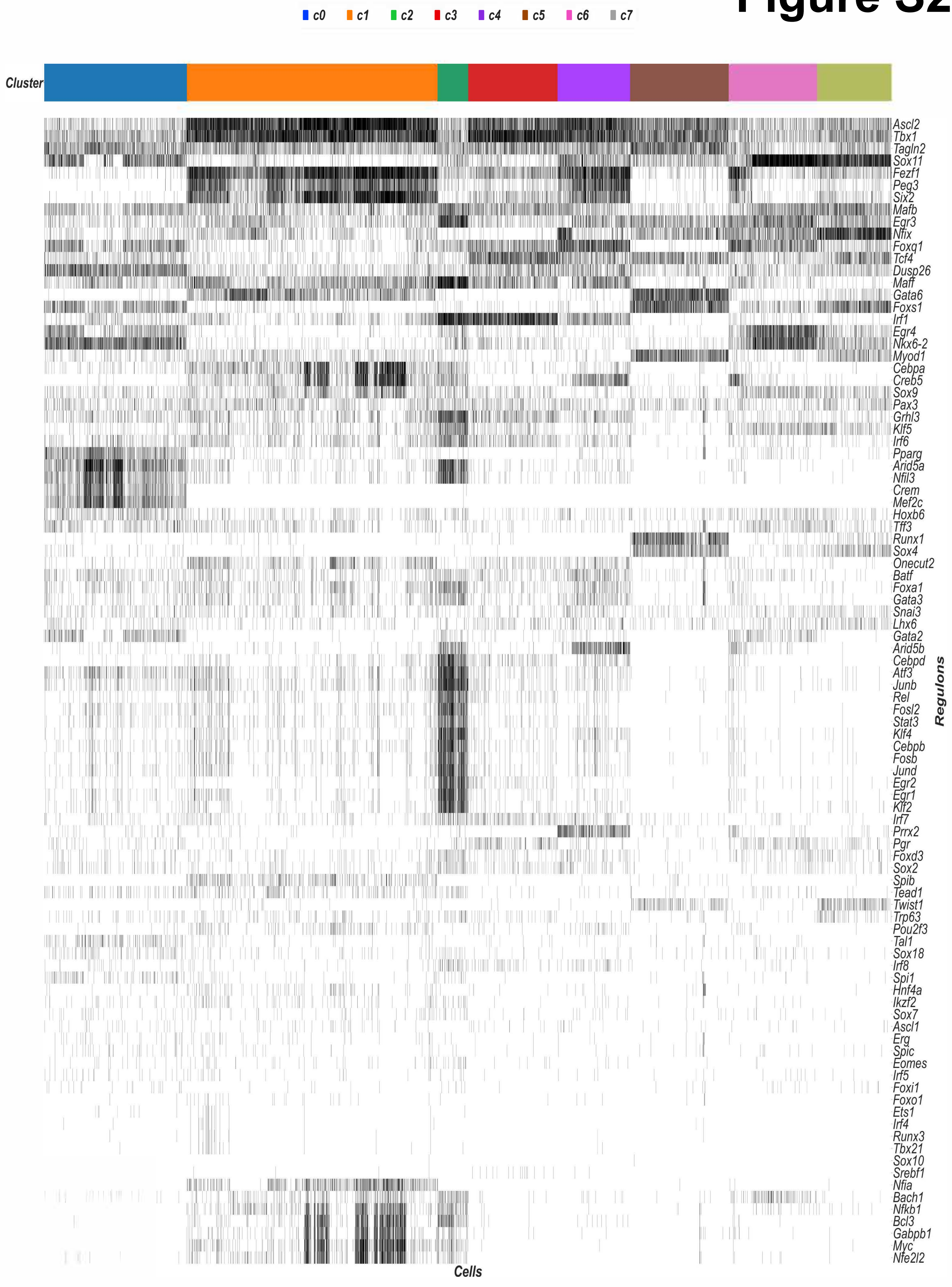
c0
c1
c2
c3
c4
c5
c6
c7
Cluster
Ascl2
Tbx1
Tagln2
Sox11
Fezf1
Peg3
Six2
Mafb
Egr3
Nfix
Foxq1
Tcf4
Dusp26
Maff
Gata6
Foxs1
Irf1
Egr4
Nkx6-2
Myod1
Cebpa
Creb5
Sox9
Pax3
Grhl3
Klf5
Irf6
Pparg
Arid5a
Nfil3
Crem
Mef2c
Hoxb6
Tff3
Runx1
Sox4
Onecut2
Batf
Foxa1
Gata3
Snai3
Lhx6
Gata2
Arid5b
Cebpd
Atf3
Junb
Rel
Fosl2
Stat3
Klf4
Cebpb
Fosb
Jund
Egr2
Egr1
Klf2
Irf7
Prrx2
Pgr
Foxd3
Sox2
Spib
Tead1
Twist1
Trp63
Pou2f3
Tal1
Sox18
Irf8
Spi1
Hnf4a
Ikzf2
Sox7
Ascl1
Erg
Spic
Eomes
Irf5
Foxi1
Foxo1
Ets1
Irf4
Runx3
Tbx21
Sox10
Srebf1
Nfia
Bach1
Nfkb1
Bcl3
Gabpb1
Myc
Nfe2l2
Regulons
Cells

### Figure S3

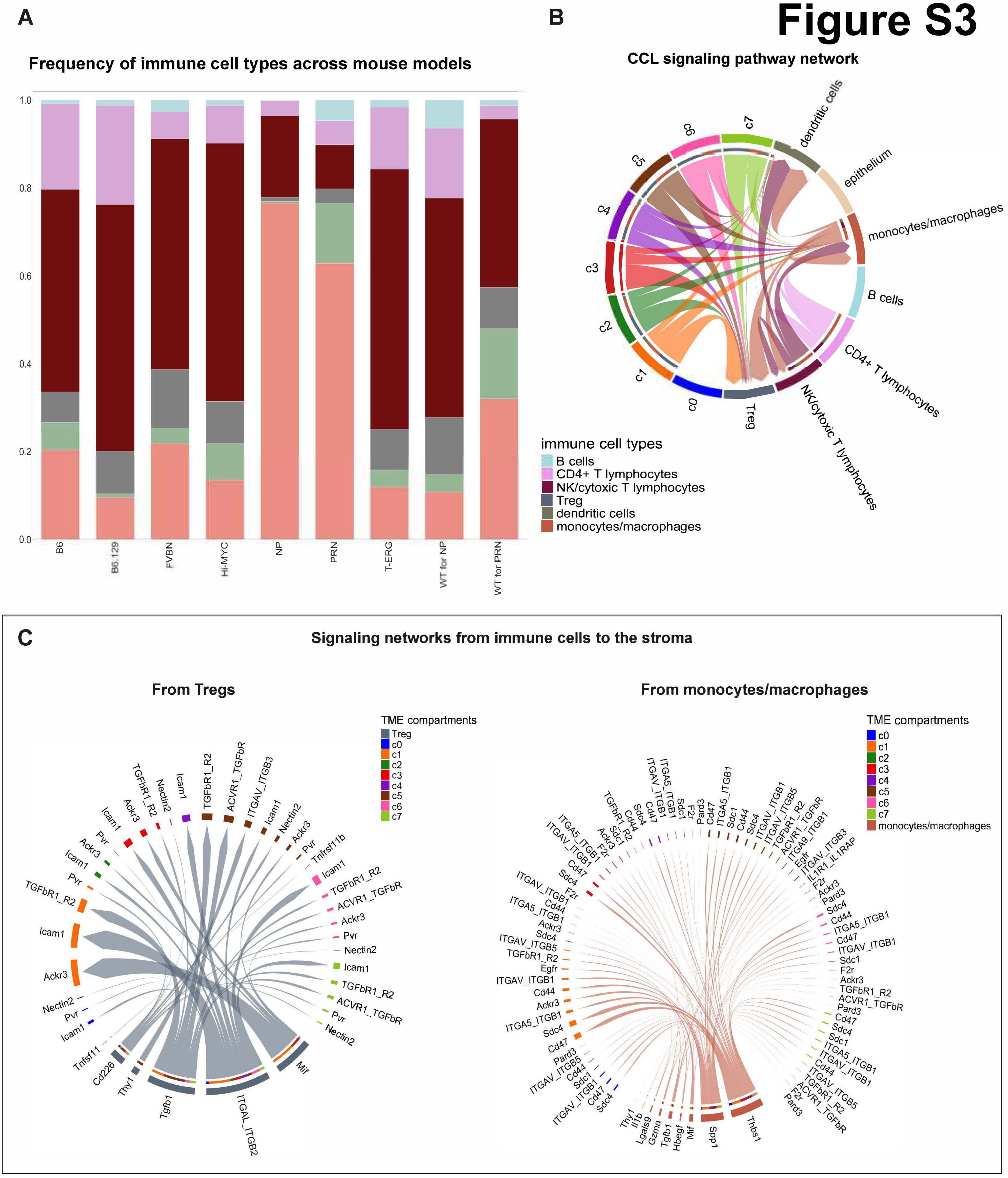

A
Frequency of immune cell types across mouse models
1.0
0.8
0.6
0.4
0.2
0.0
B6
B6.129
FVBN
Hi-MYC
NP
PRN
T-ERG
WT for NP
WT for PRN
B
CCL signaling pathway network
c7
dendritic cells
epithelium
monocytes/macrophages
B cells
CD4+ T lymphocytes
NK/cytoxic T lymphocytes
Treg
c0
c1
c2
c3
c4
c5
c6
immune cell types
B cells
CD4+ T lymphocytes
NK/cytoxic T lymphocytes
Treg
dendritic cells
monocytes/macrophages
C
Signaling networks from immune cells to the stroma
From Tregs
TME compartments
Treg
c0
c1
c2
c3
c4
c5
c6
c7
Tgfb1
ITGAL_ITGB2
Mif
Cd226
Thy1
Tnfsf11
From monocytes/macrophages
TME compartments
c0
c1
c2
c3
c4
c5
c6
c7
monocytes/macrophages
Thy1
Il1b
Lgals9
Gzma
Tgfb1
Hbegf
Mif
Spp1
Thbs1

### Figure S4

**A** **Stroma-stromal interactions**

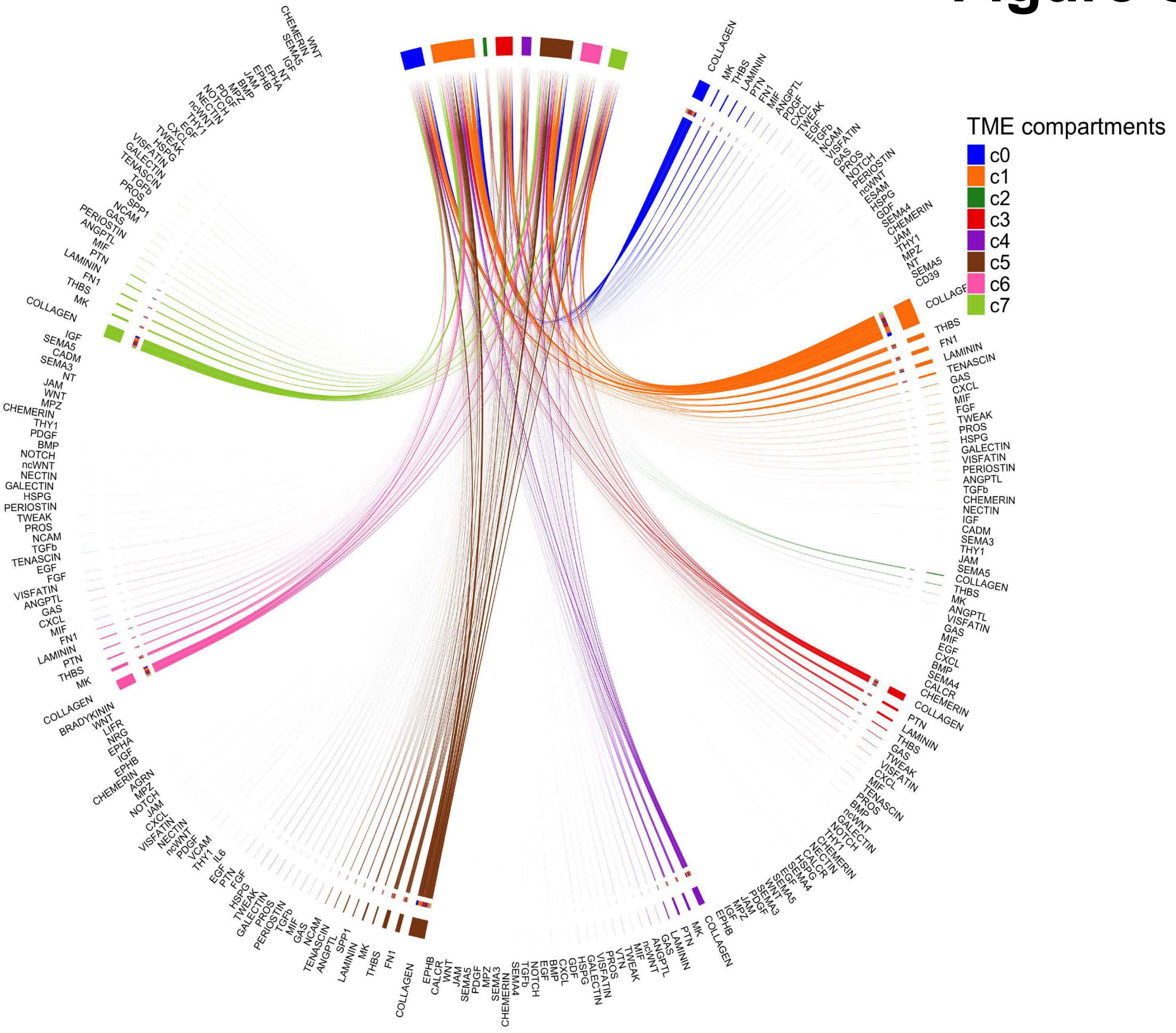


**B** **Wnt signaling within the stroma**

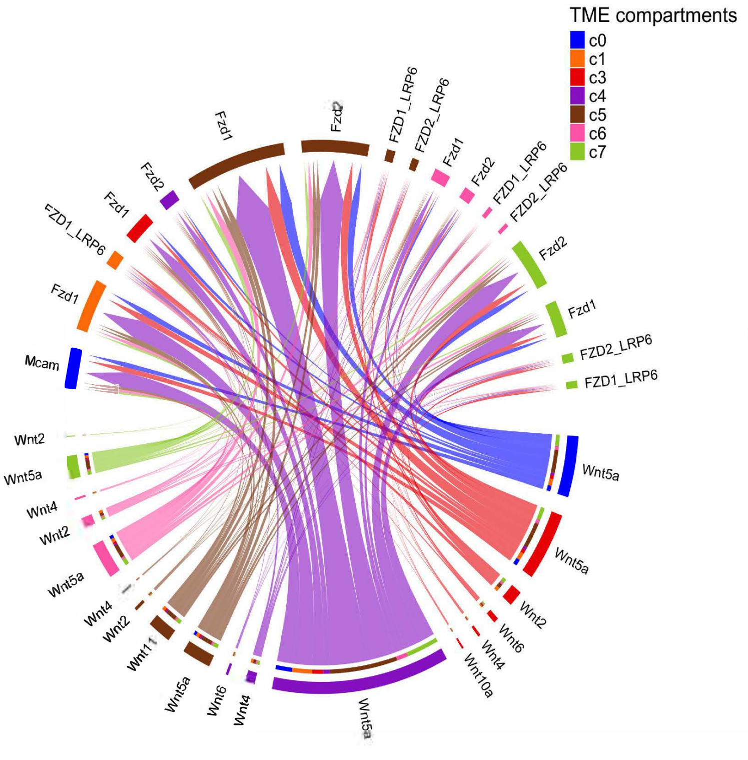


**C** **Periostin signaling within the stroma**

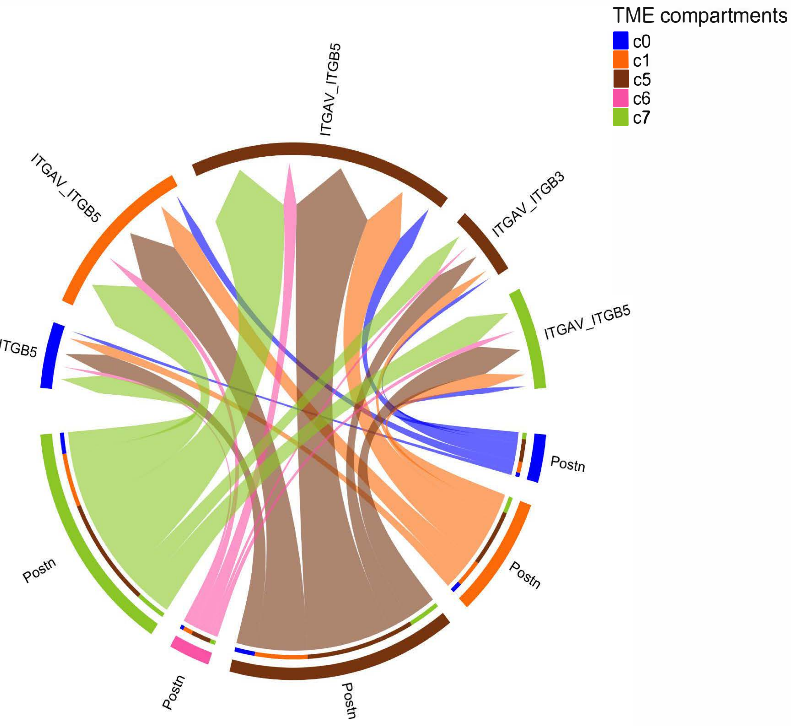

### Figure S5

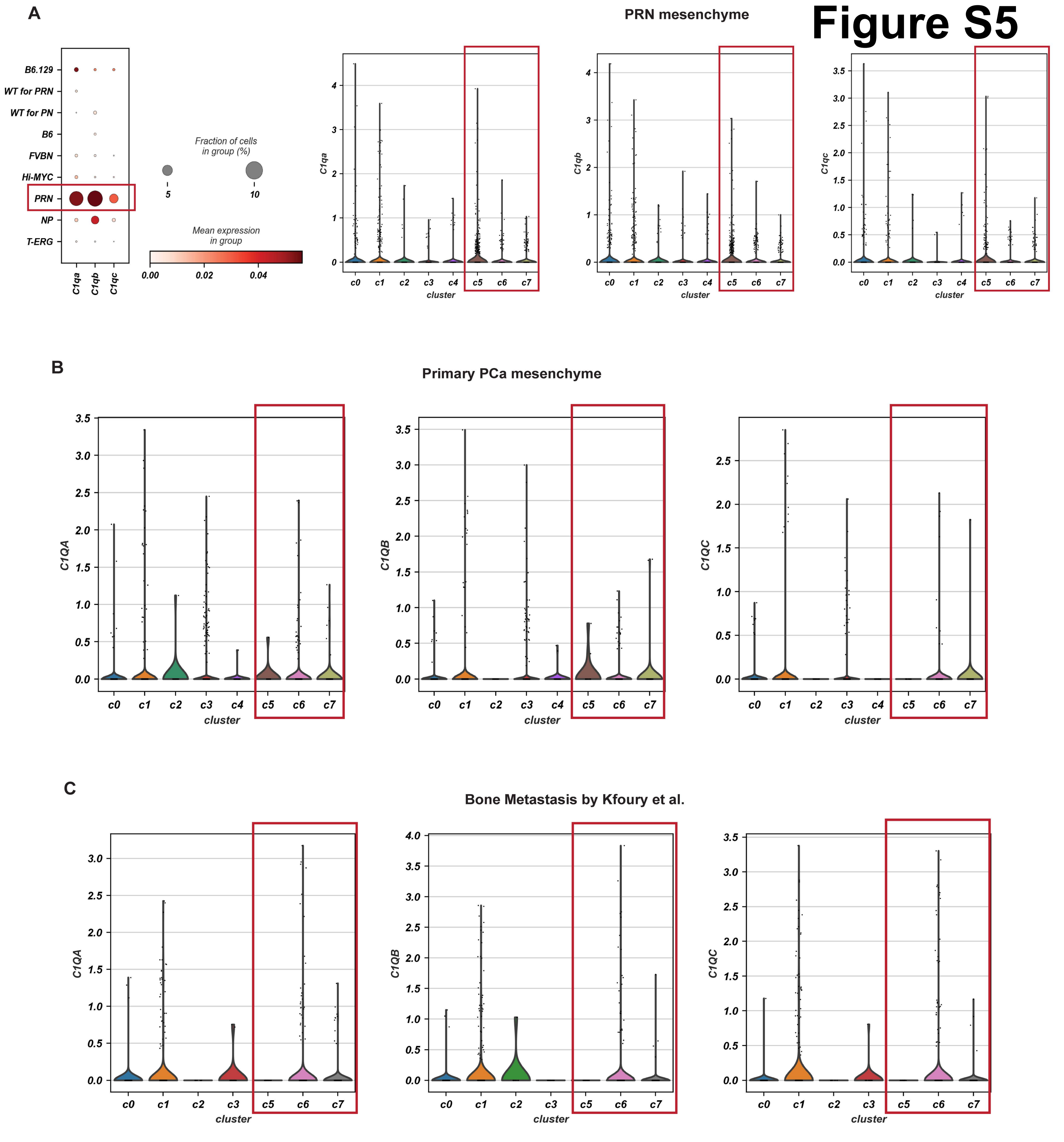
A
PRN mesenchyme
B6.129
WT for PRN
WT for PN
B6
FVBN
Hi-MYC
PRN
NP
T-ERG
C1qa
C1qb
C1qc
Fraction of cells in group (%)
5
10
Mean expression in group
0.00
0.02
0.04
c0 c1 c2 c3 c4 c5 c6 c7
cluster
B
Primary PCa mesenchyme
C1QA
C1QB
C1QC
c0 c1 c2 c3 c4 c5 c6 c7
cluster
C
Bone Metastasis by Kfoury et al.
C1QA
C1QB
C1QC
c0 c1 c2 c3 c5 c6 c7
cluster

### Figure S6

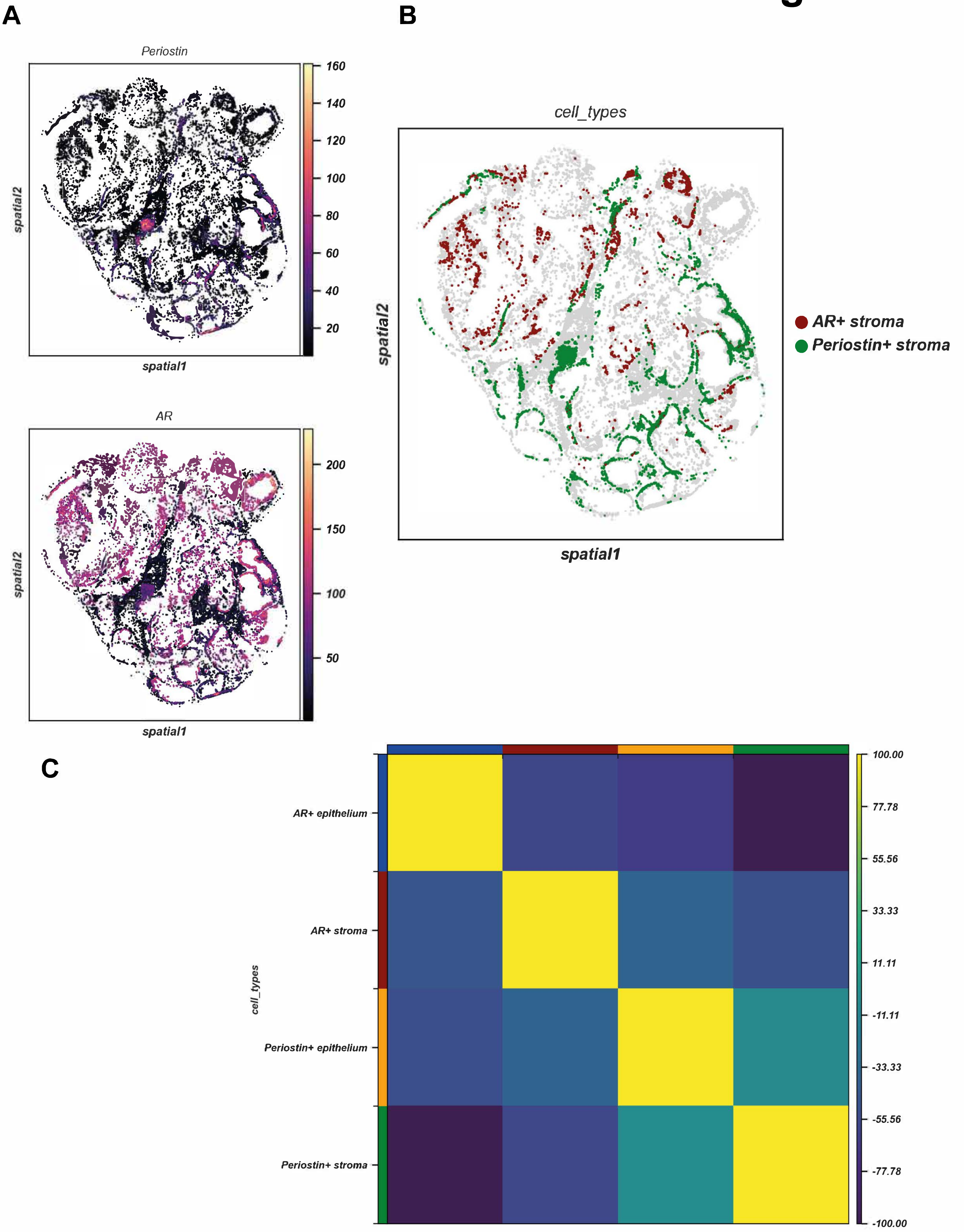
A
B
Periostin
spatial2
spatial1
160
140
120
100
80
60
40
20
AR
200
150
100
50
cell_types
AR+ stroma
Periostin+ stroma
C
AR+ epithelium
AR+ stroma
Periostin+ epithelium
Periostin+ stroma
100.00
77.78
55.56
33.33
11.11
-11.11
-33.33
-55.56
-77.78
-100.00
